# Time-varying hierarchical core voxels disclosed by *k*-core percolation on dynamic inter-voxel connectivity resting-state fMRI

**DOI:** 10.1101/2022.06.23.497413

**Authors:** Youngmin Huh, Yeon Koo Kang, Wonseok Whi, Hyekyoung Lee, Hyejin Kang, Dong Soo Lee

**Author notes:** Corresponding authors. Dong Soo Lee. Hyejin Kang.

## Abstract

*k*-core percolation on the scale-free static brain connectivity revealed hierarchical structure of inter-voxel correlations, which was successfully visualized by hyperbolic disc embedding on resting-state fMRI. In static study, flagplots and brain rendered *k*_max_-core display showed the changes of hierarchical structures of voxels belonging to functional independent components (IC). In this dynamic sliding-window study, temporal progress of hierarchical structure of voxels were investigated in individuals and in sessions of an individual. *k*_max_-core and coreness *k* values characterizing time-varying core voxels were visualized on animated stacked-histogram/flagplots and animated brain-rendered images. Resting-state fMRI of Human Connectome Project and of Kirby weekly revealed the slow progress and multiple abrupt state transitions of the voxels of coreness *k* and at the uppermost hierarchy, representing their correlative time-varying mental states in individuals and in sessions. We suggest this characteristic core voxels-IC compositions on dynamic study fingerprint the time-varying resting states of human minds.

**One Sentence Summary:** Dynamic state transitions of hierarchical functional inter-voxel connectivity implied time-varying mental states at rest on fMRI

## INTRODUCTION

Resting-state brain fMRI (rsfMRI) gives an opportunity to look into the mental state of an individual, which seems to vary along time but with a recognizable pattern or at least within a certain physiologic range at rest. Investigators assume the stationarity of mental state or its rsfMRI correlates during the entire acquisition time or the time bins that they chose. The former is said static study and the latter dynamic study of functional brain connectivity on rsfMRI (*1–7*). Dynamic study can also be done using model-based or seeds-based-brain-search ways. Model-based approach, for example, hidden Markov model (HMM), could yield time-varying states of a few ensembles of spatial principal components of brain voxels (*8–12*). Seeds-based approach was also called coactivation pattern (CAP) of searched voxels given the seeds composed of voxels of interest instead of voxels (*13–15*). Computing was the predominant concern in both approaches and choosing output states in HMM or seeds in CAP approaches was another hurdle for further comparison between or within individuals especially if the investigators were interested in individuals’ dynamic or time-varying mental states and their rsfMRI pairwise connectivity correlates.

Brain functional networks have the characteristics of an open dissipative system (*16*) with metastability (*17, 18*) resulting in the progress of states with intermittent transition (*19*), whose underlying mechanism lies in inherent composition of network nodes. Even though voxels are considered as the unit entities of brain function, current mm^3^ resolution of fMRI yield a voxel consisting of mixture of neurons of 10^5^ (*20*). Dynamic binning of time down to 1-min also let us to assume stationarity of neuronal interaction within this voxel (*21, 22*). Keeping in mind the heterogeneity of composition of voxels and unproven stationarity of time bins, we can move on to look for the hierarchical structures of voxels from both correlated and anti-correlated relations between voxels, along the pre-determined time bins with stationarity assumption. Static functional connectivity studies used long-time bins of 7 to 15 minutes with stationarity assumption. Now dynamic studies can be done using 1-min time bins to disclose whether the static and the minute time-bin dynamic analysis of rsfMRI would represent the state progress or transition of individuals’ mental states at rest (*7, 21, 22*).

We recently introduced a method to observe hierarchical structure of brain functional network, revealing the characteristic core voxels disclosed by *k*-core percolation on the adjacency matrix of static functional intervoxel connectivity acquired by the established method using <0.1 Hz rsfMRI signals (*23*). *k*-core percolation enables disclosure of functional hierarchical position of every voxel on recursive decomposition while *k* values are increased and the voxels with lower degrees are removed. *k*-max core voxels at the uppermost hierarchy are equivalent to highest collective influencers or the most vulnerable nodes. We moved this approach on to use 1-min time bins of 3 seconds shift for individuals, namely used the sliding window method to produce the consecutive 280 bins per subject. Upon completion of the analysis, the state progress/ transition of voxels representing hierarchical structures were found to vary in their compositions belonging to functional independent components (ICs). This voxel- bin approach led to devise various visualization methods, which are 1) coreness *k* flagplots, coreness *k* brain rendered images and 2) *k*_max_-core stacked histogram, bar plots or *k*_max_-core brain rendered images on animation formats.

Using these methods, intervoxel functional connectivity on rsfMRI were transformed to coreness *k* values for voxels, which meant that we could use node (voxel) instead of edge (correlation or connectivity) information for visualizing as well as interpreting hierarchical intervoxel connectivity. This also enabled us to pay attention to the intervoxel anti-correlation to yield the negative-influence networks. We put absolute values of negative correlation to the analysis pipeline to generate negative-influence networks while being cautious that the absolute value of the negative correlation represents different information from negative correlation per se. Intervoxel anti-correlation information was divided to sign (minus or plus) and value, and we could simply adopt the value information.

We read out the findings from a variety of generated plots showing coreness *k* or *k*_max_- core values for functionally connected voxels, using either correlation and anti-correlation after thresholding to confirm the scale-freeness, for inter-individuals for Human Connectome Project (HCP) and within an individual weekly (Kirby weekly), on the static (15 minutes or 7 minutes in toto, respectively for HCP or Kirby weekly) and the dynamic (280 1-min-bins with 3-sec shifts (HCP) or 166 1-min-bins with 2-sec shifts (Kirby weekly)). On the dynamic study, slow progress of k_max_-core composition and state transition were disclosed so as to manifest which ICs these hierarchically uppermost voxels belong to. We observed a variety of transition/no-transition, possible between-individual differences characterizing each individual and commonality or uniqueness of the time-varying states’ progress in individuals and sessions of an individual.

Comparing static and dynamic core characteristics of correlation as well as anti-correlation, time-varying dynamic characteristic cores were similar between individuals except for inclusive uniqueness per individuals. Distributed pattern heralded the state transition or intervened within the slow-changing states and these slow-changing states showed one to three component dominance such as visual network, default mode network plus central executive networks or others. Static product of percolation-process flagplots and stacked histogram of *k*_max_- core voxels revealed partially the convolved pattern of temporally drifting changes of hierarchical structures of voxels on resting state fMRI of individuals. Reading these animation plots/images, enabled to check the possibility/implausibility of individual/epochal (within an individual) fingerprinting to be related with mental states’ variation. We propose that *k*-core percolation on the scale-free configuration of intervoxel brain connectivity and its dynamic visualization disclosed the temporal drift of mental states and its dynamic connectivity rsfMRI correlates.

## RESULTS

### Visualization of temporal progression of resting functional hierarchical intervoxel connectivity on animation

*k*-core percolation and the visualization of voxels were reported to disclose hierarchical structures of voxels (nodes) in functional intervoxel connectivity on rsfMRI (*23*). *k*_max_-core voxels represent the uppermost voxels on these hierarchical structures, which change upon time progression spontaneously at rest during acquisition period. If we annotate *k*_max_-core voxels to functional ICs determined on group ICA, the contribution of IC subnetworks to the uppermost parts of time-varying mental states can be discovered in individuals or a repeatedly imaged single individual.

Data from HCP and Kirby weekly were put in to the analysis pipeline (Fig. 1). Briefly, intervoxel (number of voxels=5,937) correlation matrices for 280 or 166 time-binned 1-minute duration epochs of 3-seconds or 2-seconds shift, respectively for HCP or Kirby weekly data, were put in for display of degree distribution to find the inverse linearity on the log-log plots for the determination of threshold, to confirm the scale-freeness of degree distribution with average 88.5% of the entire voxels included after thresholding (*24*). In HCP data, threshold was found to be 0.65 and in Kirby weekly, 0.8 to meet the criteria of as many voxels included but with confirmed scale-free degree distribution.

**Fig. 1.**
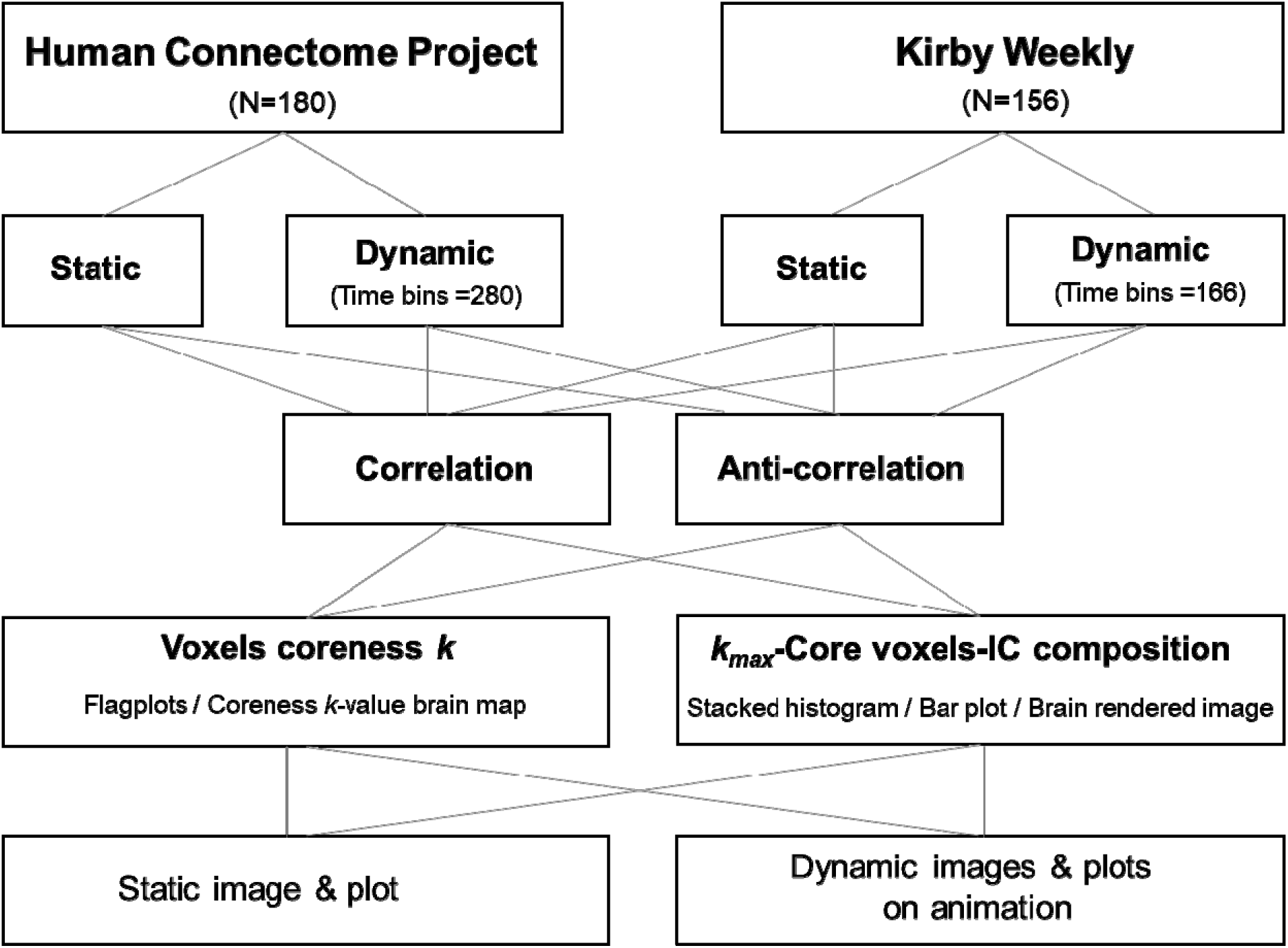
Analysis and Visualization. Data from Human Connectome Project were for analysis for individuals. Data from Kirby weekly for sessions from an individual. Static plots for voxel coreness *k* values and *k*_max_-core IC composition were compared with those on animation. IC: independent components.

Adjacency matrices made with these thresholds were put in to *k*-core percolation (*23, 25*), which produced 5,937 x 280 percolated matrix for *k* values. These *k* values of each voxel were rearranged according to their belonging to ICs (Fig. S1) (default mode network (DMN), salience network (SN), dorsal attention network (DAN), central executive network (CEN), sensorimotor network (SMN), auditory network (AN), visual network (VN) and unclassified (UNC)) to be shown as 280 flagplots on animation in case of HCP data (Fig. 1, Movie S1). Kirby weekly data were processed with 5,937 x 166 percolated matrix, otherwise the same as HCP (Fig. 1, Movie S2).

The visualization of *k* values of voxels on *k*-core percolation was with flagplots on animation (Fig. 2A, Movie S1 & S2) and coreness *k* value maps on brain axial slices on animation (Movie S3). Flagplots on animation were showing the characteristic time-varying changes of coreness *k* values upon node-pruning by *k*-core percolation. Flagplots showed the pruning pattern of each IC-belonged voxels upon percolation and the façade of each IC-voxels at the end of percolation showed the number of *k*_max_-core voxels, and the length of flags represented the depth of hierarchical structures of IC-voxels, for example, the percolation steps 200 for AN, 225 for SN, and 250 for DMN/CEN/VN. Coreness *k* value brain maps were composed of axial/sagittal/coronal and 31 layers of transaxial brain-rendered images. These coreness *k* values derived from the static and the dynamic studies have matrix forms (Fig. S2) and are ready for further analysis such as variational autoencoder (VAE) or statistical parametric mapping (SPM) for anomaly detection.

**Fig. 2.**
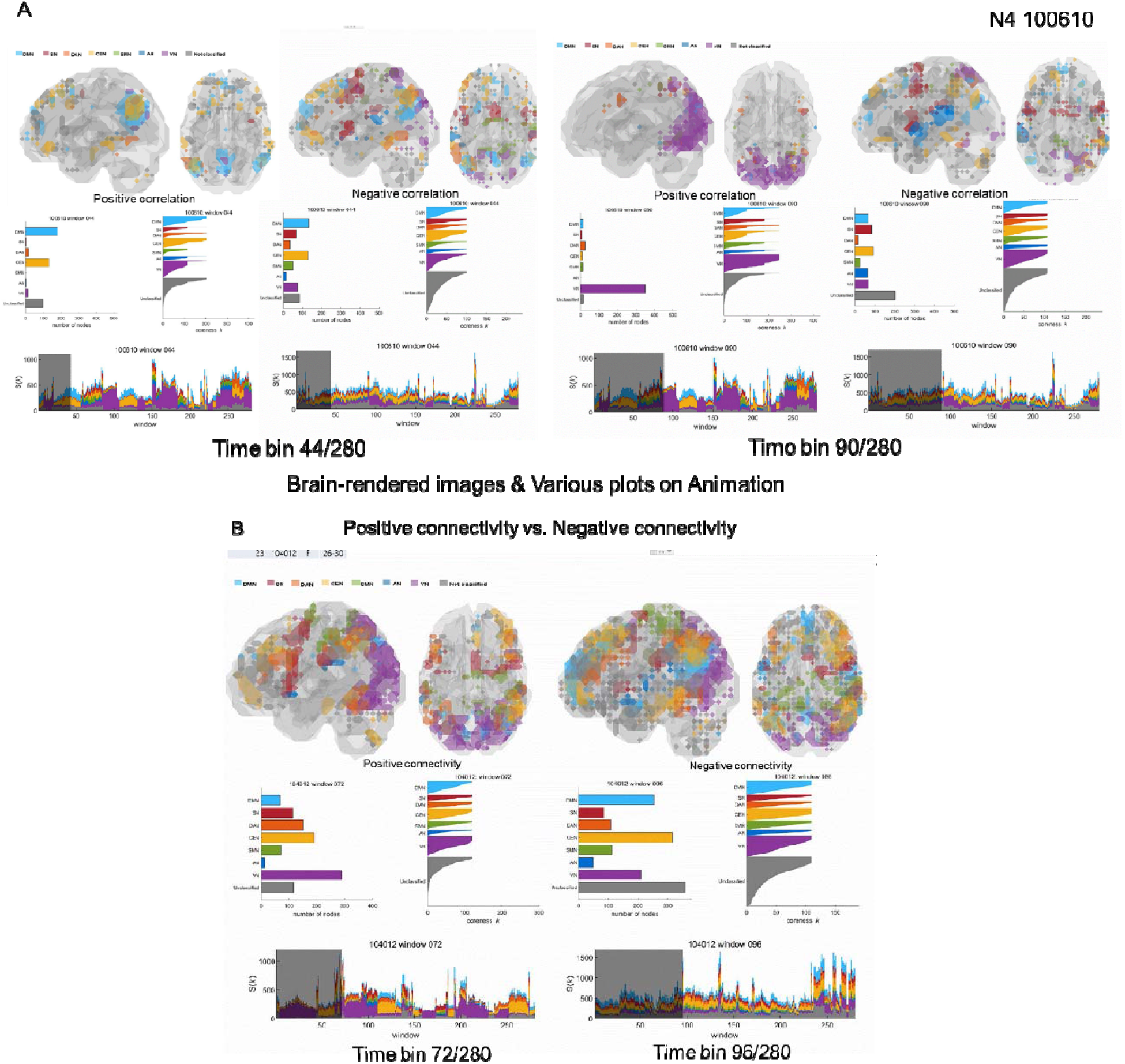
Visualization outputs for static and dynamic analysis. Images and plots on animation (See Movie S4 & S5) were shown for two snapshots at the time bins of different epochs for an individual and another. A) At 44/280 time bin on the left, positive connectivity on the left, negative connectivity on the right. At 90/280 time bin on the right, too. Negative connectivity results were similar between 44/280 and 90/280 bins, but positive connectivity were varying from DMN/CEN dominant *k*_max_- core voxels to visual dominant (See Movie S4) . B) In another individual, on the left, at the time bin of 72/280, distributed pattern of *k*_max_-core voxels from positive connectivity was different from the distributed one of negative connectivity on the right (See Movie S5).

The uppermost voxels were designated as *k*_max_-core voxels (*23*), and plotted in three ways, 1) bar plots showing numbers of *k*_max_-core voxels on animation, 2) stacked histograms DMN/SN/DAN/CEN/SMN/AN/VN/UNC *k*_max_-core voxels colored with cyan-rainbow colors- gray on animation, and 3) axial and sagittal brain-rendered images containing *k*_max_-core voxels colored the same way to their belonging to ICs and on animation (Fig. 2A, Movie S4; Fig. 2B, Movie S5). Stacked histogram and bar plots ignored the location within eight subneworks (DMN/SN/DAN/CEN/SMN/AN/VN/UNC) and right/left-sidedness of *k*_max_-core voxels. Thus, further examination of brain-rendered axial images disclosed the asymmetry of *k*_max_-core voxels (Fig. S3, Movie S6). States, transitions, pulses (or trains) were counted on visual read-outs on animated images for the voxels belonging to ICs and summed for the individuals in toto and for the sessions of an individual from Kirby weekly.

### Visual assessment of *k*_max_-core voxels showing hierarchical structure of functional connectivity in individuals on static study

Static study of functional intervoxel positive connectivity revealed variations in dominant (VN or DMN) or distributed patterns on the *k*_max_-core voxels-ICs composition analysis (Fig. 3A, B). Among 180 individuals, 109 showed distributed pattern with voxels of VN, DMN, SMN the most prevalent, 32 showed co-dominant voxels of DMN/VN, and 19 DMN 18 VN the dominants. 25 showed co-dominant voxels of UNC (19 of them along with DMN). 75 of 180 showed cerebellar *k*_max_-core voxels mostly belonging to UNC and rarely CEN/DMN within the cerebellum (Fig. 3C).

**Fig. 3.**
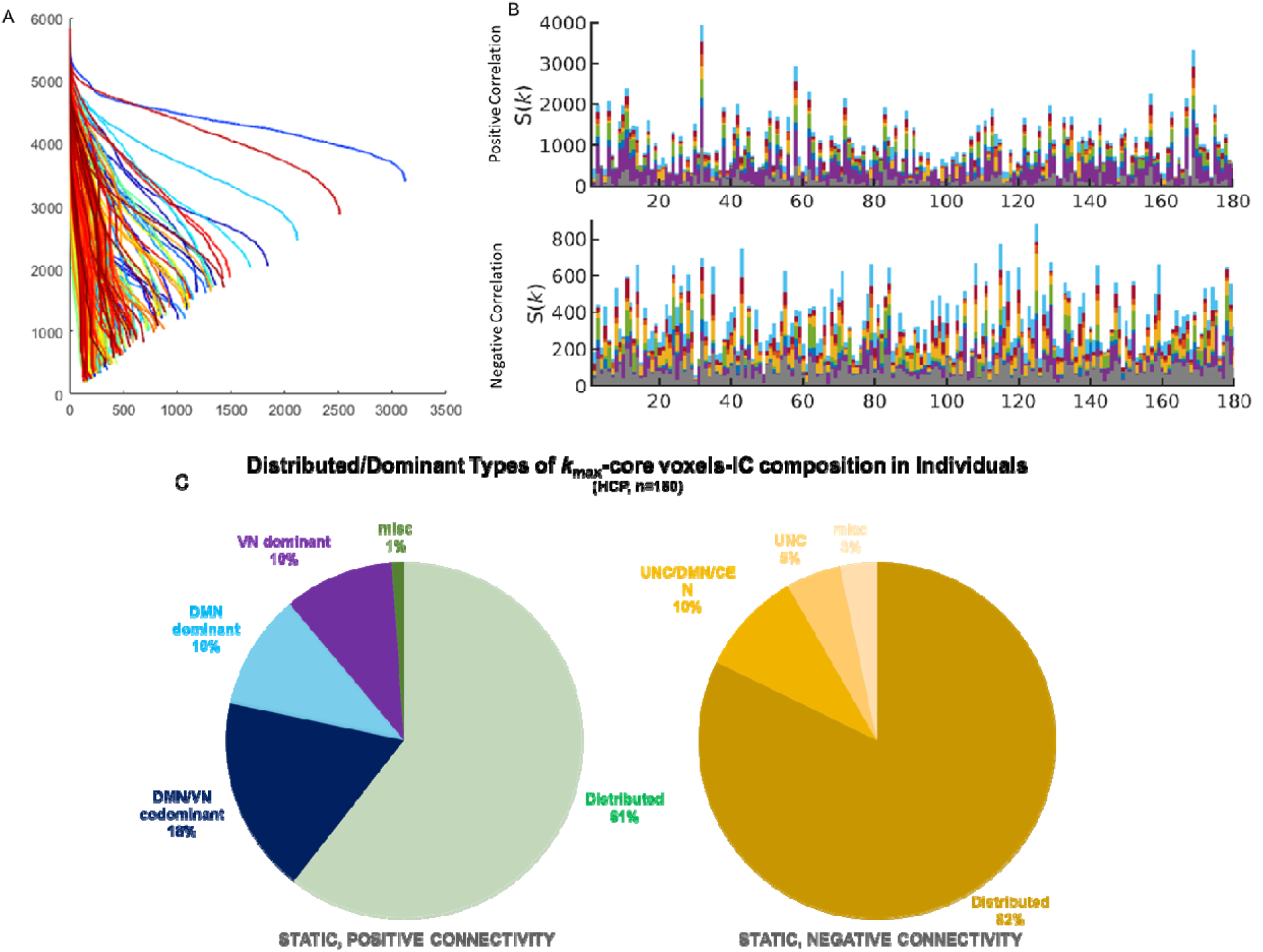
Stacked histogram plots of k_max_-core voxels of individuals for positive and negative correlations on static study. A) *k*-core percolation and their varied descent of coreness *k* values of the entire voxels, B) HCP database, static analysis of individuals (n=180) showed varied size *k*_max_-core voxels and IC composition, C) Dominance or distributed pattern of *k*_max_- core voxels-IC composition between individuals. More than half of the individuals were distributed, and DMN or VN dominants were the remainders in positive connectivity and much more were distributed in negative connectivity.

Static study of negative connectivity (its absolute value) revealed similar and also unique patterns on the IC composition of the *k*_max_-core voxels. 148 individuals showed distributed pattern, however, rarely with UNC, SMN, CEN dominance. 26 showed co-dominant voxels of UNC (17 of them along with DMN/CEN). The other 10 showed dominant voxels of other ICs or combinations thereof. 108 of 180 individuals showed cerebellar *k*_max_-core voxels similarly to positive connectivity (Fig. 3C).

### Visual assessment of progress/transitions of hierarchical structure of functional connectivity in individuals on dynamic study

Dynamic study of positive connectivity revealed much more diverse pattern of dominance and the distributed one as well as their time-varying progresses, whose dominant voxels belonging to VN, DMN/CEN ICs showed the slow progress and abrupt change of the voxels-ICs composition (Fig. 4A). State transition was visually defined as an abrupt voxels-IC composition changes to another composition. Slow progress was visually traceable ones of the same voxels-IC composition for some while (several bins to even the entire 280 bins).

**Fig. 4.**
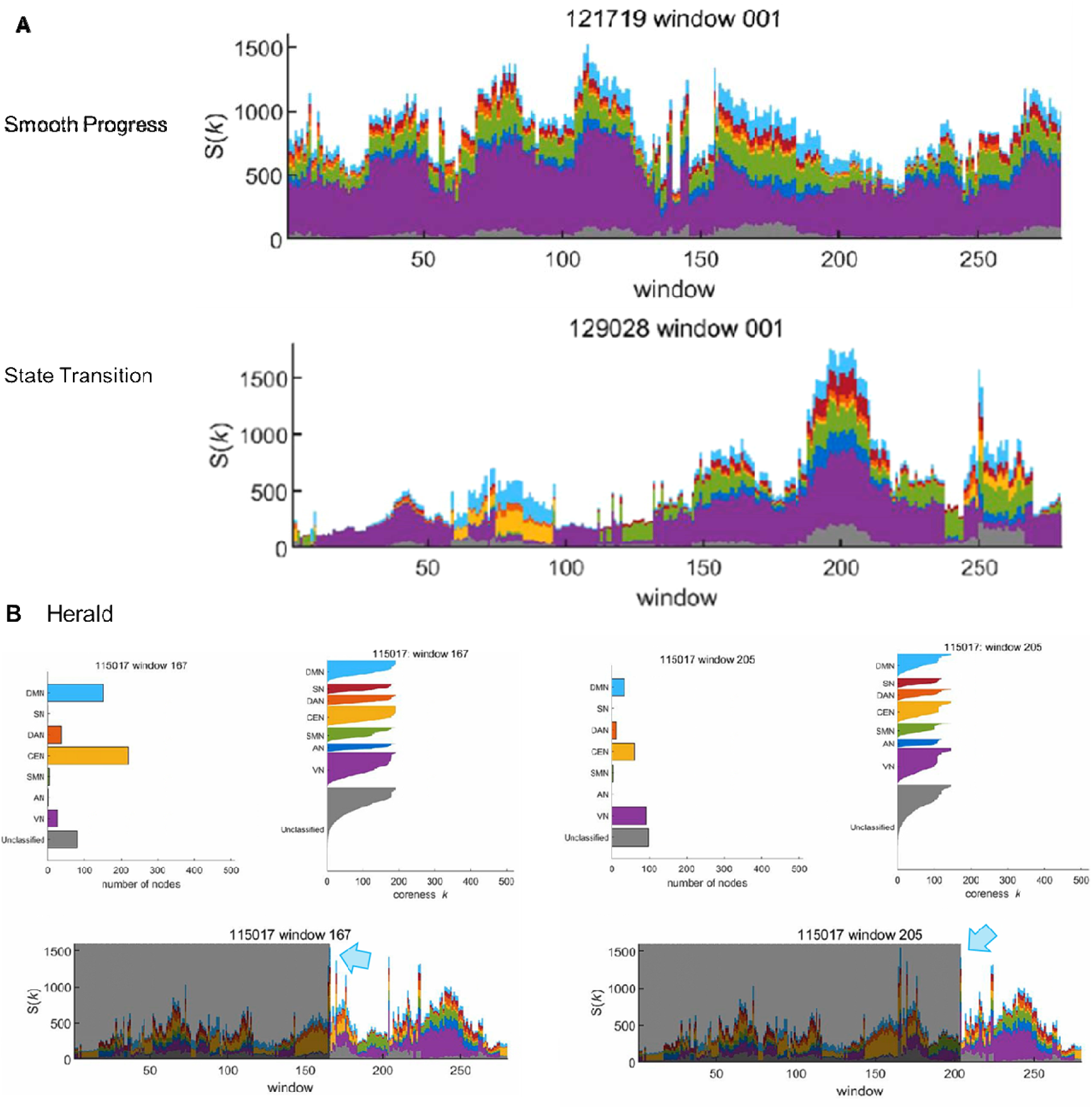
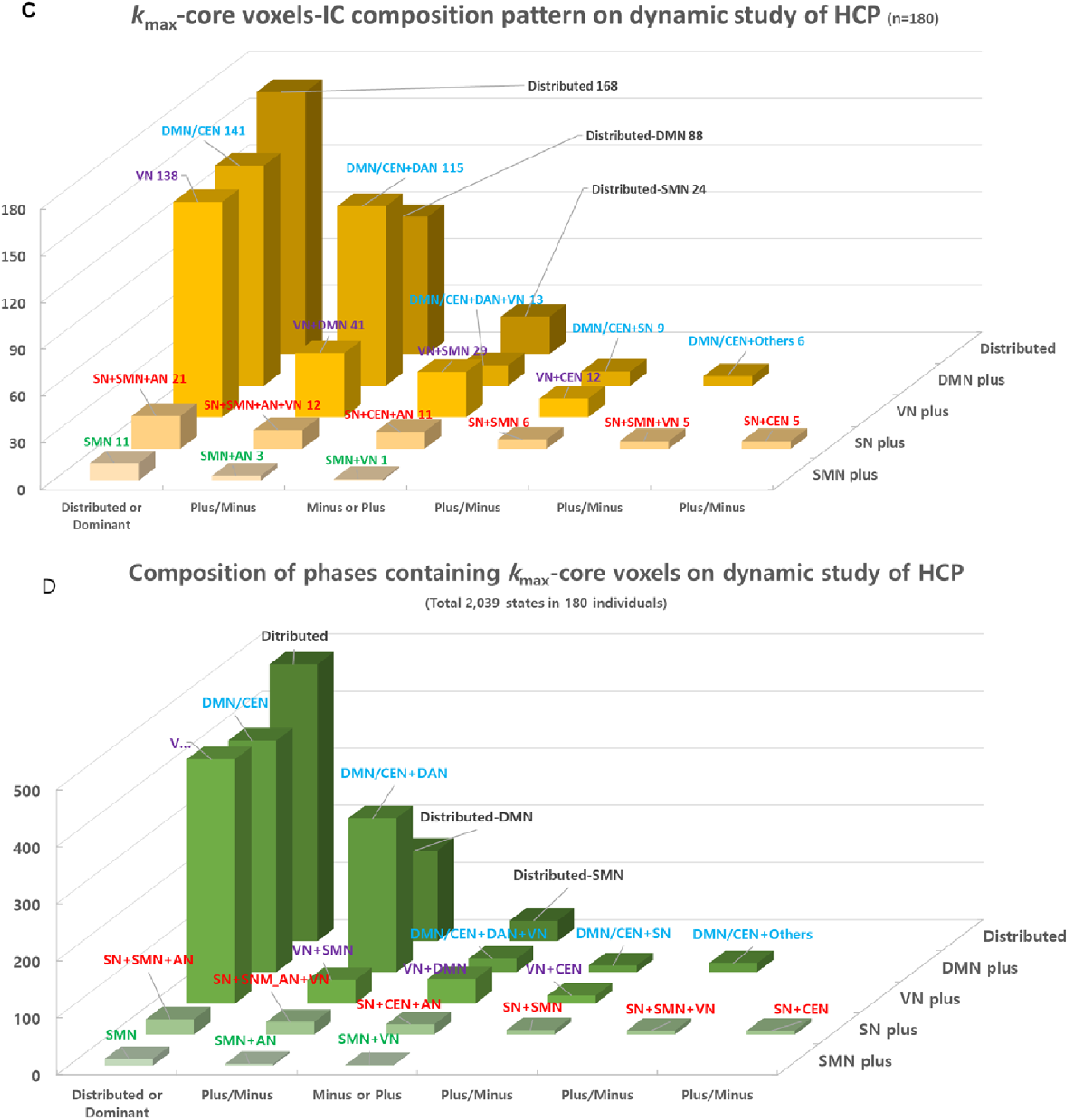
Time-varying dominance/distributed k_max_-core voxels-IC composition on dynamic study. A) An individual’s dynamic study showed rare state transition but slow progress VN- dominant and also the distributed pattern of *k*_max_-core voxels (upper). Another individual’s dynamic study showed state transition from VN-dominant to DMN/CEN/VN co-dominant and to DMN/CEN-co-dominant and then again back to VN-dominant. This pattern of changing vertical strips while taking turns to occupy the uppermost position of the IC-hierarchy competition was called state transition. B) Two separate single time-bins were indicated, one successful to herald state transition, and another not. Herald bins are supposed to represent brain voxels’ inherent motion to take the shifts of voxels-IC composition for the next period. C) Among 180 individuals, how many showed one or more dominance/distributed pattern along 280 time-bins were represented as bars. Distributed and the similar, DMN/CEN and the analogues, VN and additives were major patterns while minor ones were SN/SMN and the related, or SMN and the relates ones. D) Total 2,039 states were determined on visual examination, which also revealed similar composition in terms of fraction of the total in the entire population of 180 with their 280 time-bins total (180 x 280).

Reproducibility of visual determination of state transition in individuals was with the correlation of 0.88 for repeated read-outs by one operator among authors, transition numbers being ranged from 2 to 21 per individuals (Fig. S4). Pulses of duration one to several bins showed up intermittently, which counted 0 to 18 times per individual (mean 6.4+3.6). These pulses heralded the following state transition (Fig. 4B).

*k*_max_-core voxels-IC compositions were read out to reveal the prevalence of distributed or DMN/CEN, VN, SN/SMN or SMN dominant patterns of recognizable states within each individual and along all the individuals. Among the 180 individuals from HCP, the individuals were most prevalent who could be annotated as showing the distributed, DMN-short distributed (distributed lacking only DMN) or SMN-short distributed types. DMN/CEN or DMN/CEN plus (DAN, DAN/VN, SN or others) were the next prevalent types. SN/SNM and its plus types or SN and its plus types followed (Fig. 4C). Among the total 2,039 annotated states along the entire 180 individuals, distributed and DMN or SMN-short distributed types were also the most prevalent, DMN/CEN and its plus types and VN and its plus types followed. All the states of distributed, DMN/CEN, VN and other SN/SMN or SN were observed without bias all over the individuals.

No prominent characteristics of whether a special type of *k*_max_-core voxels-IC composition was observed in each individual and thus the commonality of *k*_max_-core voxels-IC composition prevailed in all the studied individuals. Within each individual, any tendency leaning to certain specific combinations of *k*_max_-core voxels-IC composition during the moment of rsfMRI acquisition could not be found but cannot be totally excluded either (Fig. 4D).

Dynamic study of negative connectivity was unique in its revelation of eloquent commonality between individuals. Unlike positive connectivity, temporal progress of states were not so explicit, in other words, the voxels belonging to ICs almost always showed up along the temporal progress of *k*_max_-core voxels composition, sometimes withdrawing from the uppermost position but quickly returning (Movie S1 & S2). Unclassified voxels, which we annotated for convenience sake, sustained to be present, all over the cerebrum and also cerebellum.

Localization or dominance was not so noticeable in most of the individuals, meaning distributed pattern prevailed along all the individuals and along all the time of observation by rsfMRI (Movie S4-S6). In few cases, *k*_max_-core voxels-IC composition of dynamic negative connectivity mimicked the pattern of dynamic positive connectivity.

Flagplots revealed distinctive pattern of positive and negative connectivity; in the positive connectivity, the total number of *k*_max_-core voxels tended to increase proportionally with the length of *k*-core percolation, which meant the depth (or height) of hierarchical structure of voxels, whereas in the negative connectivity, the length of *k*-core percolation changed over time independently from the total number of *k*_max_-core voxels, which implied there were inherent rhythm to make the global hierarchical structure fluctuate/undulate along the time lapses (Fig. 5, Movie S7). Sum of all the voxels in IC-designated flags was equivalent to summed voxels’ coreness k values on degree-distribution plots (Fig. S5)

**Fig. 5.**
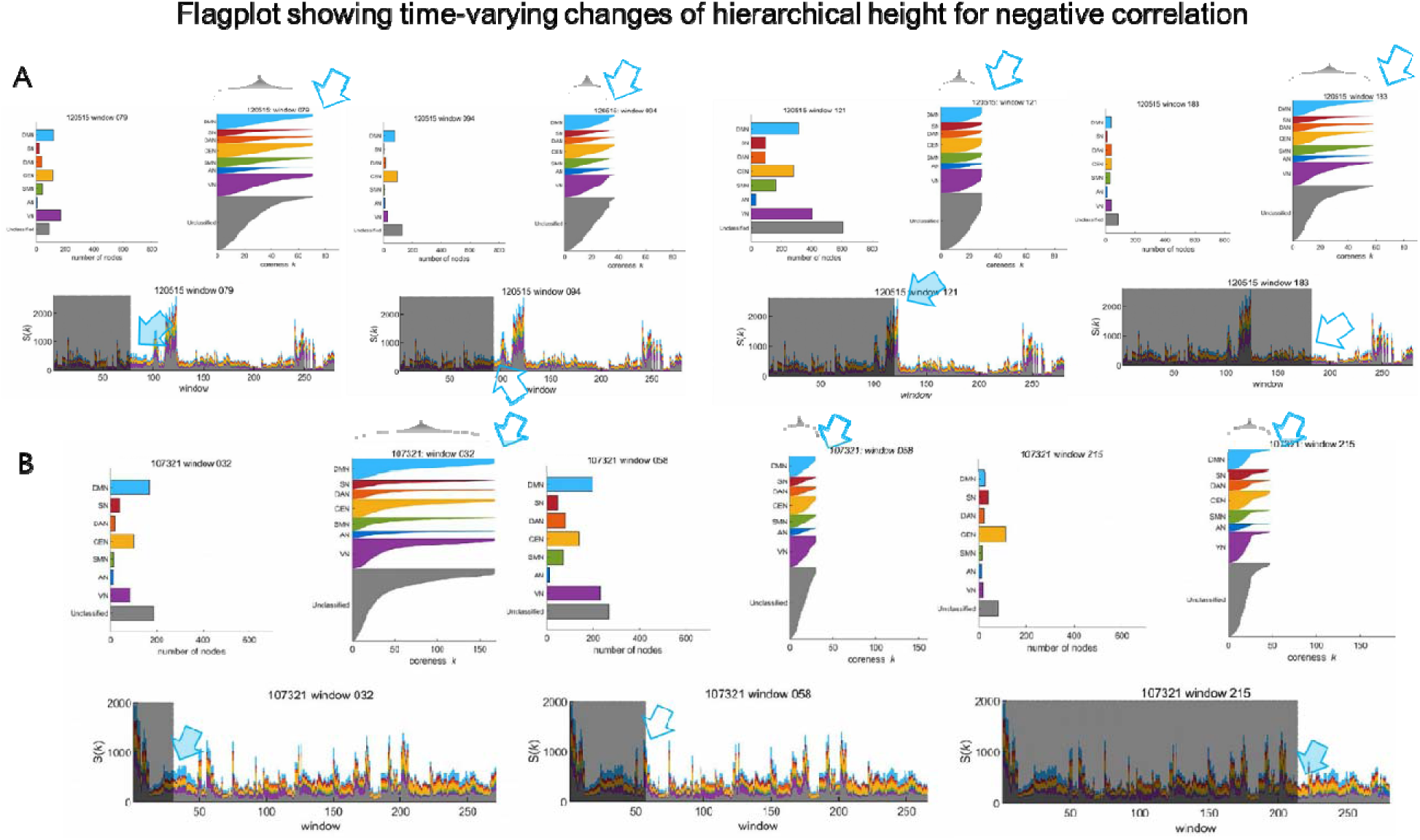
Flagplots showing the height/depth of hierarchical structure of each IC- voxels along k-core percolation See Movie S7). A) In an individual, flagplots on animation made by dynamic analysis using intervoxel negative correlation showed lengthening and shortening of the flags belonging to each IC, disclosing the time-varying changes of hierarchical depth. *k*_max_-core voxels were just presented as the width at the end of the flags. Pulse, tall eruption of one or more bins, was not associated with the lengthening of the flags but rather shortening thereof. B) In another individual, despite the plain progress with just the ripples of bins, the length of the flags changed on their own, implying the flag-lengthening related increase of voxels hierarchy which was supposed to represent inherent rhythmic ensemble motion of independent multiple voxels in the brain.

Brain-rendered images of *k*_max_-core voxels of dynamic study could reveal the sameness or the difference (despite the same belonging to ICs) of participating voxels along time. Details could be visualized and described and the asymmetry of the left or the right hemisphere could very well be commented on brain-rendered images. In positive connectivity, right/left asymmetric participation of voxels to *k*_max_-core were found in 40 individuals (Fig S2, Movie S6), however, in negative connectivity, asymmetry of *k*_max_-core voxels-IC composition was rare though not absent. On brain-rendered images, we also could discover the participation of cerebellar voxels more prevalently in negative connectivity, where voxels belong to UNC most and flickering/blinking of cerebellar DMN and CEN along time.

### Comparison of static hierarchical structure of and dynamic temporal progress of hierarchical structures of functional connectivity

*k*_max_-core voxels-IC composition of static study, represented as bar plots, stacked bar plots and flagplots (5,937x1 vector) were compared with temporal progress of dynamic study, represented as animated bar plots, animated stacked histogram plots, animated flagplots (5,937x 280 voxel vector x time bin matrices) (Fig. 6, Fig. S2). Not in positive connectivity, but in negative connectivity, the fraction of *k*_max_-core voxels-IC composition of the animated 280 stacked histogram plots (the time sum) of the dynamic study was proportional with the fraction of *k*_max_-core voxels-IC composition of stacked bar plot of the static study (Movie S8-S10).

**Fig. 6.**
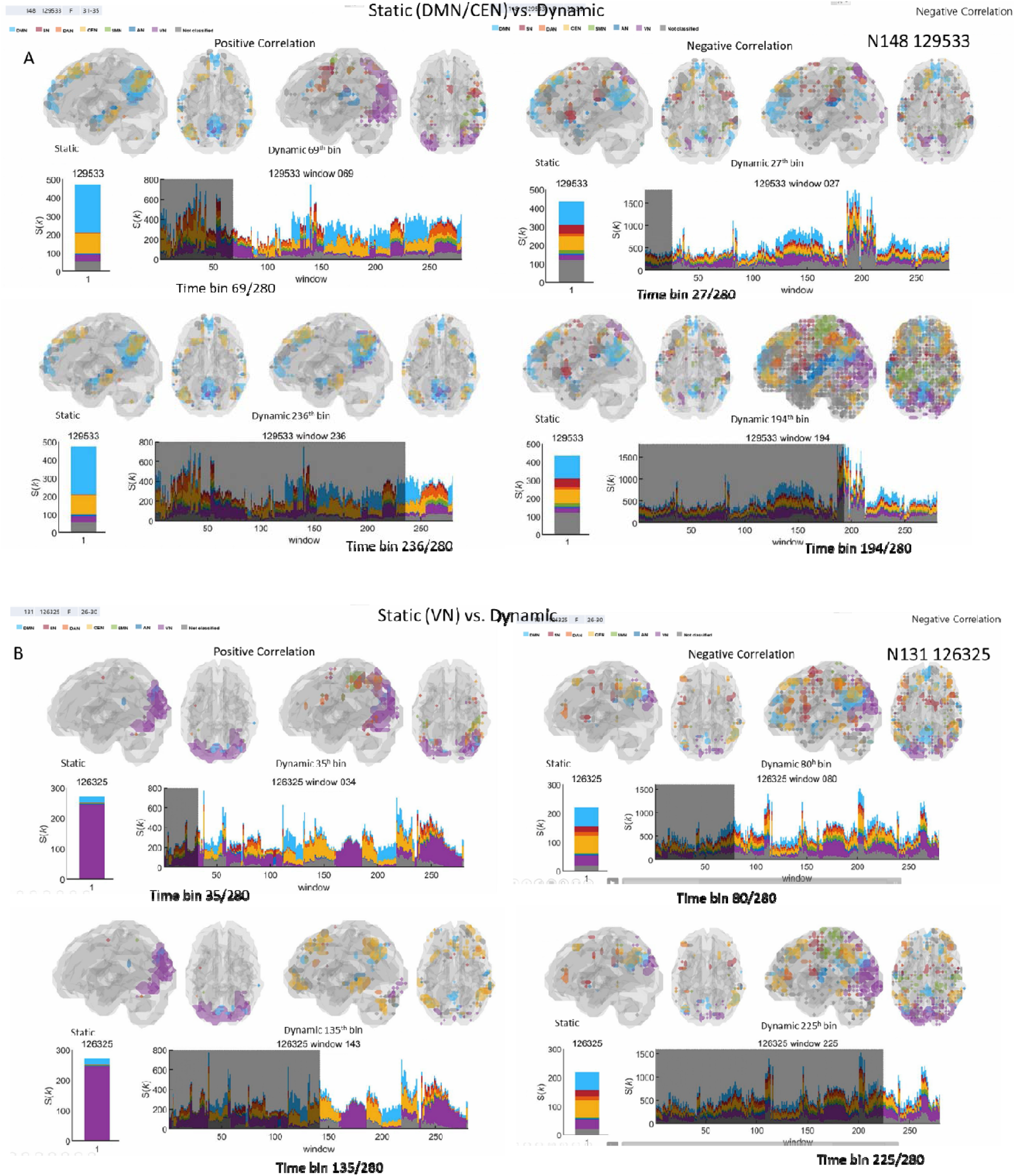
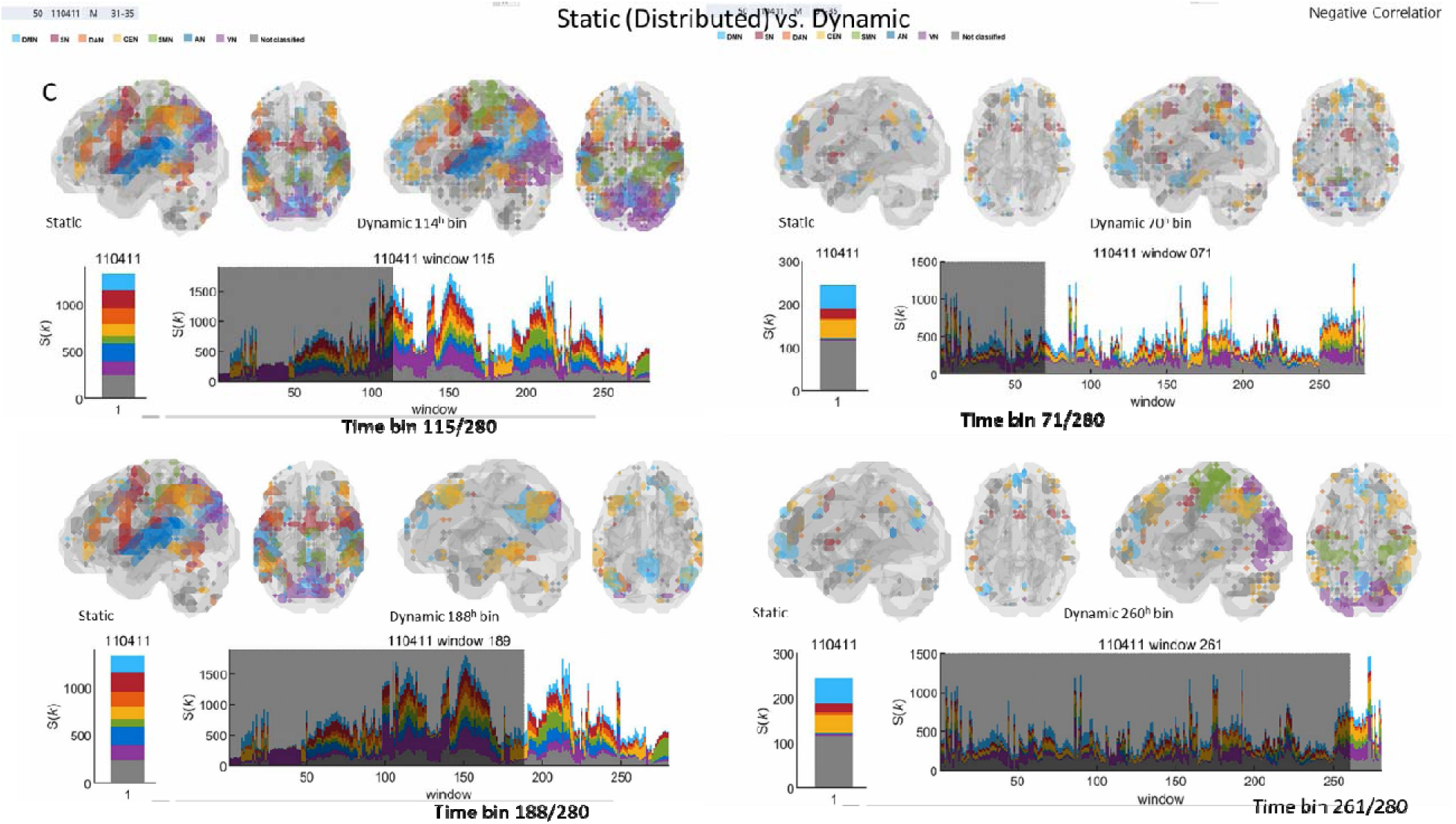
Comparison of static plots/images with dynamic plots/images on animation. A) DMN/CEN pattern on static, but both VN dominant and DMN/CEN are observed on dynamic for positive connectivity for positive connectivity on the left side (See Movie S8). B) VN pattern on static, but additional DMN/CEN are also there for positive connectivity (See Movie S9). C) Distributed pattern on static, and also distributed major but with minor DMN/CEN on dynamic for positive connectivity (See Movie S10). For negative connectivity pictured on the right sides, ambiguous between individuals.

In positive connectivity, neither qualitative (kinds of ICs, pattern of compositions, size of voxels to IC, their progress or transition etc.) nor quantitative (number of voxels belonging to each IC and their sum over time per IC and their sum again) comparison revealed any similarity (Fig. 6). Simply saying, in positive connectivity unlike negative connectivity, the bar plot or the stacked bar plot did not enable the prediction of temporal changes of animated stacked histogram plots or animated bar plots. Interestingly, the reverse was also the case, meaning that observation of the entire dynamic study did not lead to any proper guess of the stacked bar plots of static study.

On brain-rendered images, gross pattern of flickering *k*_max_-core voxels belonging to ICs were similar between static study and dynamic study in negative connectivity (Fig. 6, Movie S8- S10). In positive connectivity, this was not the case, meaning dynamic study revealed slow progress and abrupt state transitions of *k*_max_-core voxels-IC composition in their own way and static study revealed its own representative *k*_max_-core voxels-IC composition per individual.

Nevertheless, in positive connectivity, fractional prevalence of states among all the 2,039 states of entire 180 individuals were similar to that of static study (Fig. 3C left pie plot, Fig. 4D, Fig. 6). In both static and dynamic study, from the most frequent to the least were the distributed pattern, DMN/CEN codominant pattern, VN dominant pattern and others.

Individuals showing the distributed pattern in static study, showed DMN/CEN codominant or VN dominant patterns as well as all the other patterns of *k*_max_-core voxels-IC composition. Individuals showing DMN/CEN codominant pattern in static study showed distributed or VN dominant patterns and other *k*_max_-core voxels-IC composition patterns in dynamic study. It was also the case with the individuals showing VN-dominant pattern or those showing other minor *k*_max_-core voxels-IC composition patterns.

### Diverse and varying hierarchical structure of functional intervoxel connectivity in repeated weekly sessions in an individual

In an individual studied repeatedly on rsfMRI 156 times weekly in Kirby weekly study, in static study of positive connectivity, DMN was dominant in 130 sessions (59 with CEN, 38 with DAN), VN was dominant in 68 sessions, while this VN dominant was with DMN in 57. The distributed type was few. Voxels in the cerebellum participated *k*_max_-core in 10 sessions for static positive connectivity. In static study of negative connectivity, the distributed was dominant in 147 sessions unlike that in positive connectivity. The other 8 sessions were with DMN (with CEN and/or DAN). Voxels in the cerebellum participated *k*_max_-core in 99 sessions among 156 for static negative connectivity (Fig. 7A).

**Fig. 7.**
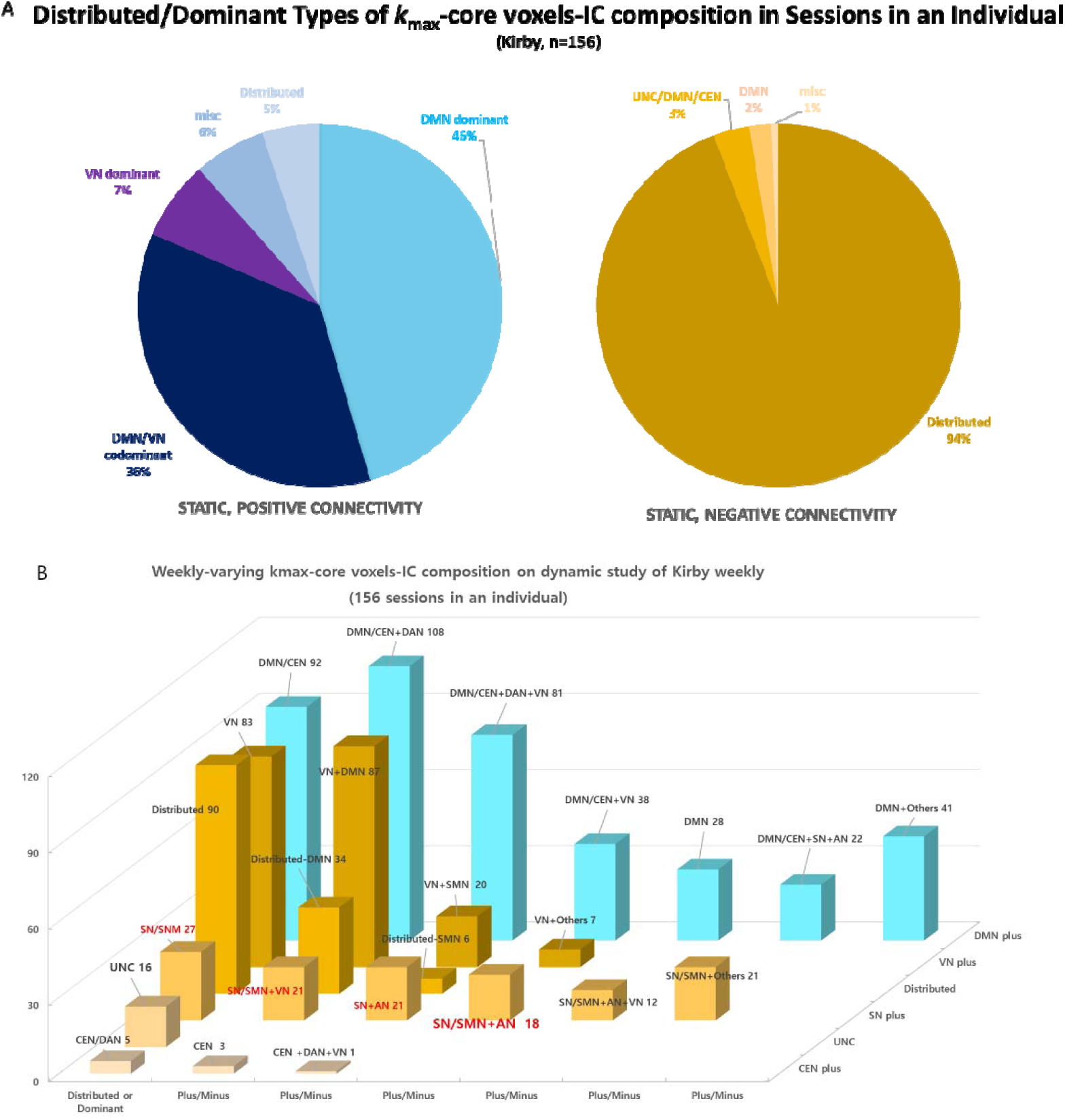
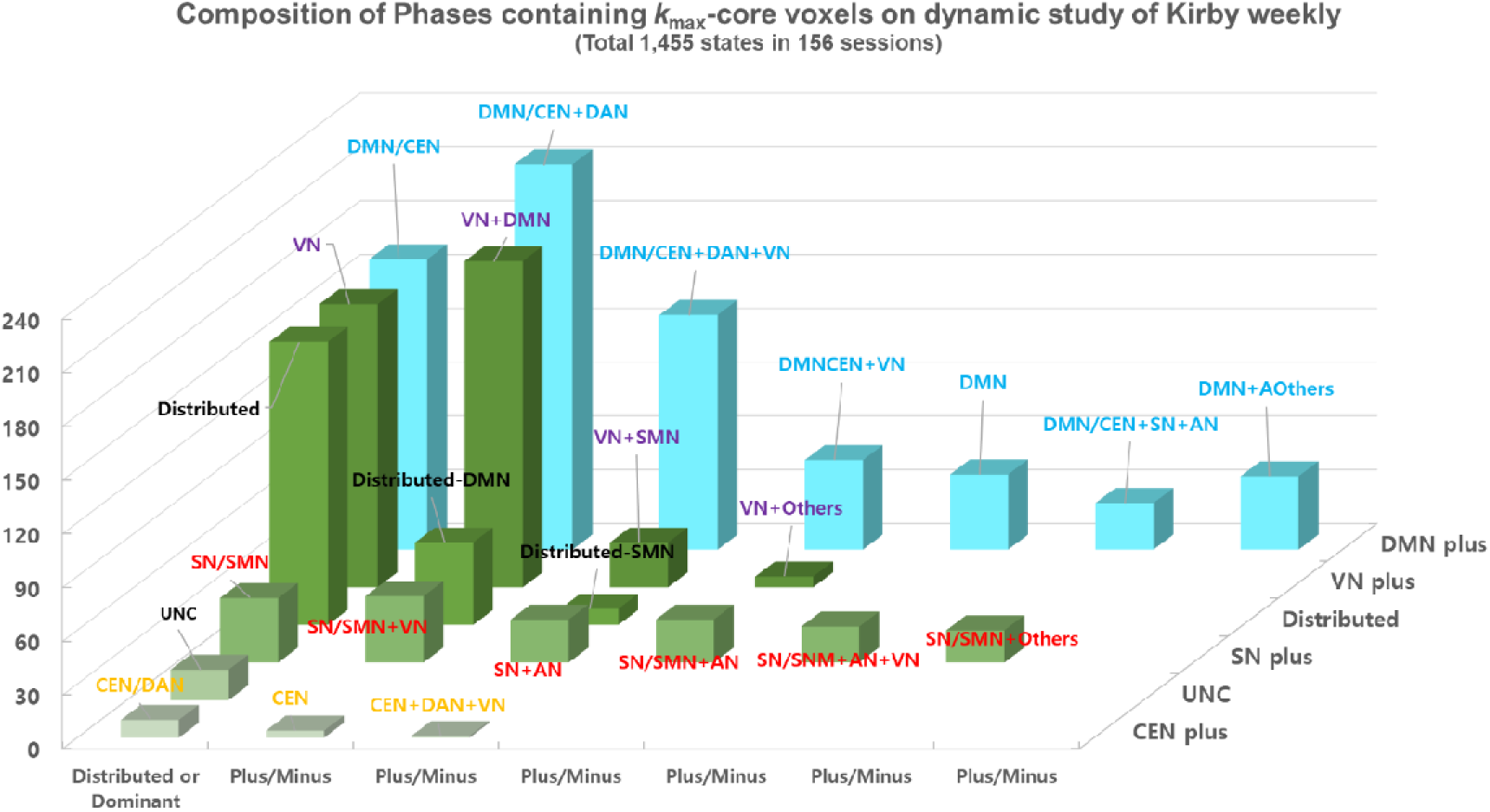
*k*_max_-core voxels-IC composition for positive and negative correlations on static and dynamic study among sessions of an individual. A) On static study, dominance or distributed pattern of *k*_max_-core voxels-IC composition between 156 sessions of an individual of Kirby weekly study. A half of sessions were DMN/CEN co-dominant, one third DMN/VN co- dominant, and remainders were VN dominant or distributed or miscellaneous patterns in positive connectivity. In negative connectivity in contrast, almost all were distributed pattern. B) Among 156 sessions on dynamic study, DMN/CEN dominant and the analogues dominated, VN dominant and additives followed and distributed pattern was the next. SN/SNM and UNC and CEN patterns were recognized rarely. C) Total 1,455 states over the sum of 156 sessions were determined on visual examination, which also revealed similar composition in terms of fraction of the total in the entire session of 156 with their 166 time-bins total.

In positive connectivity of dynamic study, reproducibility of designation of state transition per session showed the correlation of 0.87 between repeated read-outs (state transition per session 4-20; mean=10 S.D.= 3.0) (Fig. S4). Dominant states among 156 repeated examinations were in the order of prevalence, 1) DMN/CEN co-dominant type, 2) VN-dominant and distributed types and then 3) the minor SN/SMN or CEN types (Fig. 7B). States defined visually were numbered 1,455 for the entire 156 session (4-17 per individual in one examination; mean=9.3 S.D.=2.7). Weekly pattern of *k*_max_-core voxels-IC composition among 156 sessions varied from the most prevalent DMN/CEN codominant, the second most VN-dominant, the third distributed and then the other minor SN/SMN co-dominant or rare CEN dominant types (Fig. 7C). Pulses showed up intermittently, which counted 0 to 15 times per session (mean 4.8±2.6). These pulses also heralded the following state transition. Voxels in the cerebellum participated *k*_max_-core in 10 sessions. Voxels in the cerebellum participated *k*_max_-core in 151 sessions for dynamic positive connectivity.

In negative connectivity dynamic study, like individuals’ study of HCP database, in an individual, the voxels of almost every IC persisted along the temporal progress of *k*_max_-core (Fig 8A, 8B). Unclassified voxels continued to be present, over the cerebrum and the cerebellum.

**Fig. 8.**
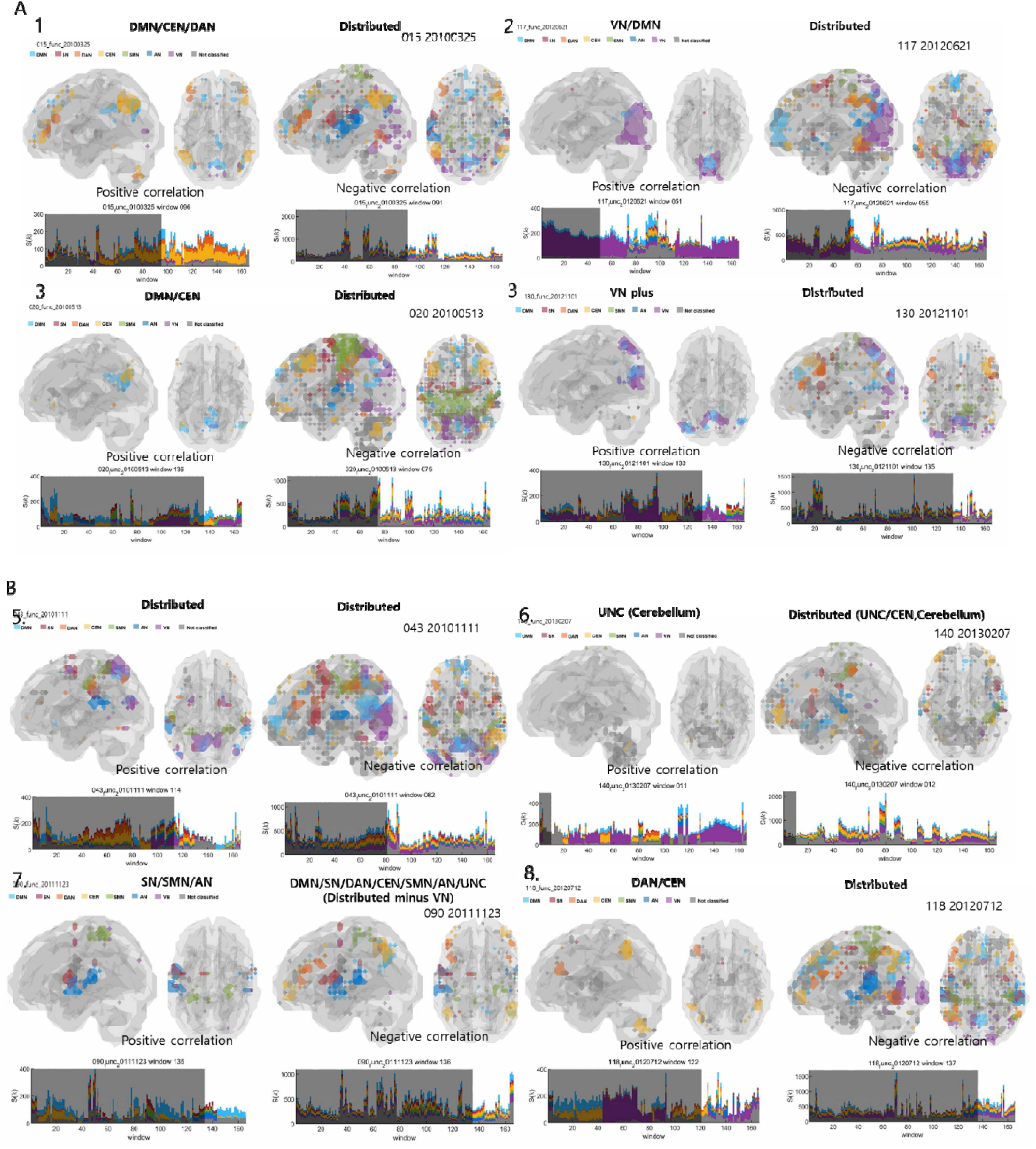
Comparison between *k*_max_-core voxels-IC composition of positive and negative correlation analysis among multiple sessions in an individual. Multiple occasions of IC-voxels dominance/distributed patterns were compared in positive and negative correlation analyses for representative sessions. A) DMN/CEN/DAN-distributed, DMN/CEN-distributed, VN/DMN-distributed and VN/DMN-distributed combinations on positive-negative correlation analysis (See Movie S11), B) Distributed-distributed, SN/SNM/AN-Distributed without VN, UNC (mainly cerebellum)-distributed, DAN/CEN-distributed combinations. Especially, UNC- distributed combined pattern was unique in that unclassified including many cerebellar voxels were the uppermost voxels on *k*-core percolation (See Movie S12).

Distributed pattern prevailed along all the sessions (Movie S11, S12). In few cases, *k*_max_-core voxels-IC composition of dynamic negative connectivity mimicked the pattern of dynamic positive connectivity.

Flagplots of positive connectivity of entire sessions suggested the length of flags could be considered to represent depth of hierarchical structure of voxels of each IC and thus the entire voxels. In positive connectivity, length increased to yield larger numbers of *k*_max_-core voxels rather proportionally, but in negative connectivity, the length of flags varied along time independently from the numbers of *k*_max_-core voxels. The latter findings on negative connectivity was comprehensible in that the depth of functional hierarchical structure might ride its own rhythm of temporal changes, whose voxels were working as ensembles. In contrast, the proportional flickering on positive connectivity might represent regional/globally concordant movements to ride up the hierarchical structure by the propensity of *k*_max_-core voxels to the uppermost hierarchy. In few sessions, brain-rendered images allowed the observation of the asymmetry of voxels’ participation of *k*_max_-core. Flickering/blinking of voxels disclosed the fact that the voxels belonging to the ICs took turns to be in charge of *k*_max_-core, as in between-individuals study of HCP.

*k*_max_-core voxels-IC composition of static study, only in negative connectivity, regardless of sessions, the fraction of *k*_max_-core voxels-IC composition of the animated 166 stacked histogram plots (the time sum) of the dynamic study was proportional to the fraction of *k*_max_-core voxels-IC composition of stacked bar plot of the static study. Unlike negative connectivity, in positive connectivity, static study results did not enable us to predict dynamic study results and vice versa.

*k*_max_-core voxels-IC composition of negative connectivity, this Kirby weekly (an) individual’s characteristics was ambiguous meaning not characteristic from the population of 180 individuals of HCP. In contrast, *k*_max_-core voxels-IC composition of positive connectivity, this Kirby weekly individual’s pattern was rather unique in that this person used more frequently DMN/CEN co-dominant voxels at the uppermost hierarchy of functional connectivity, then VN and the less the distributed. This person’s uniqueness suggested the plausibility of fingerprinting, if given the findings from the entire 156 sessions, and if an individual were examined thoroughly against 180 or more HCP individuals.

### Quantitative assessment of time-varying hierarchical structure of functional connectivity on dynamic study

In our previous investigation (*23*) and in static analysis of HCP data of this investigation, we counted the number of individuals showing VN-dominant (50% of the voxels belonging to VN), DMN/CEN co-dominant or distributed among 30 or 180 individuals, respectively.

Prevalence of distributed/dominance types was presented. On negative connectivity analysis, 82% of 180 individuals had distributed type, among the remaining 32, DMN, UNC and CEN were the dominants (Fig. 3C). Pie or percent plots summarized the static study results.

On dynamic study, states and state transitions were determined on visual analysis. Though sometimes the state determinations are obscure, all the individuals could easily be determined to have any of the distributed, DMN/CEN co-dominance or VN dominance of *k*_max_- core voxels along the 280 time-bins (Fig. 4C). Consequent sum-up of the prevalence of these patterns yielded the prevalence for the total 180 individuals (Fig. 4D). Similarly, for dynamic study of 156 sessions of Kirby weekly, counting the state transitions, number of states, number of dominant types within session (Fig. 7B) and over the entire sessions (Fig. 7C) yielded concordant results.

Flagplots of static study of previous investigation (*23*) enabled us to follow the changes of the length (depth) of hierarchical structure of each IC-affiliated voxels on *k*-core percolation. On dynamic study, time-varying changes could be followed for the voxels per each IC on animation as well (Fig. 5, Movie S7). Time-varying changes of the flag lengths per ICs explain the functional dynamics of collective entities (belonging to ICs) made of voxels as multiple independent generators.

Dynamics of *k*_max_-core voxels at the uppermost position of *k*-core percolation were presented on animation as bar plots, stacked histogram plots and brain-rendered axial/sagittal images. Coreness *k-*values of voxels representing the position of each voxel in the hierarchical structure in time-matrix format (Fig. S2) was rendered as brain-rendered images upon the tomographic transaxial templates of MRI on animation (Fig. S6, Movie S3).

### Commonality and uniqueness of ***k***-core voxels composition and their transition along time in individuals and in an individual in dynamic studies

Commonality of the *k***-**core voxels-IC composition was easily recognized on the read-outs of dynamic animation format of flagplots, which showed the pattern of slimming of the width of flags along the percolation process per each ICs. This percolation-related slimming of flags of coreness *k*-values per voxels were put on the time-varying pattern detection on dynamic animation frame. Human operator could easily recognize the pattern of changes, including slow progress of the same distributed/dominant *k*_max_-core voxels-ICs composition and the abrupt transition of these compositions. A variety of progress and transition were observed between individuals and between sessions in positive connectivity analyses, but the progress and composition was more similar between individuals and sessions in negative connectivity analyses. This was found despite the differences of epoch timing, total time of acquisition and other parameters between HCP and Kirby weekly.

Uniqueness of *k*-core voxels of any parameter on state progresses and transitions, their number, pattern, or any peculiarity was looked for especially on stacked histograms on animation or brain-rendered images on animation, *k*_max_-core voxels-ICs composition on animation and coreness *k-*value brain map and flagplots on animation. Each individual could be characterized mostly on stacked histogram and brain-rendered images on animation. Each session could be also characterized over the entire sessions to suggest any peculiarity for certain combination of *k*_max_-core voxels-IC compositions for this individual, such as SN-AN dominant or CEN dominant patterns.

## DISCUSSION

Dynamic analysis of rsfMRI using *k*-core percolation disclosed the hierarchical structure of functional intervoxel connectivity and time-varying changes of core voxels according to the coreness-*k* values of voxels and *k*_max_-core voxels-IC compositions. One-minute time bins of individuals and repeated sessions of an individual, on this dynamic study, yielded coreness-*k* values of the entire brain while revealing the voxels at the uppermost hierarchy on *k*-core percolation, tandemly over hundreds of overlapping time bins. On animation, especially stacked histogram plots of the *k*_max_-core voxels, distributed/dominant types of core voxels/IC composition showed smooth temporal progresses or abrupt state transitions during each acquisition period. Coreness-*k* values of voxels on flagplots on animation, depth of the hierarchical structure on the percolation showed global fluctuation for both positive and negative (anti-correlation based) connectivity analyses. The width of each IC at the max showing the number of remaining voxels at the max are equivalent to the number of voxels on the bar plots or brain-rendered images on animation. Read-outs of these animated plots and images enabled counting the states/ state transitions/ voxels-composition to characterize individuals and each session of an individual to propose for or against possibility of characterization of an individual among many individuals or a session (or sessions) of an individual among many sessions.

Previously observed uniqueness per subject of IC-*k*_max_-core composition in static study were observed in dynamic study too and its comparison was done with static IC-*k*_max_-core composition. Each distributed/dominant composition pattern of static study in individuals was observed, as a commonality, on each individual’s dynamic study as well as on repeated sessions of an individual. We suggest that dynamic analysis of rsfMRI using *k*-core percolation with its visualization tools on animation opened a window through which investigators look into the time-varying changes of mental states at resting state.

For dynamic study of functional connectivity, definition of spatiotemporal quanta is prerequisite for further analysis. Sliding windows had frequently been used but with a variety of protocols. Spatial units ranged from voxels (1 mm^3^) to voxels-of-interest (VOIs; hundreds or more of voxels). In HMM analysis, initially voxels were used but for definition of states, these voxels were aggregated to make a few to dozen state units of interpretation consisting of thousands or more of voxels (*8, 9*). In CAP analysis, though the search space is composed of voxels in the entire brain, seeds were made of thousands of voxels (*13–15*). In any cases, aggregation of voxels mandates the assumption that the VOIs are spatially homogeneous in terms of temporal propagation of fMRI signals, which is infrequently and inadvertently the case. Alternative is using voxels themselves as input and sustaining to use them throughout the analysis, which we adopted as a strategy to avoid arbitrariness and heterogeneity in selecting VOIs. Of course, the inherent heterogeneity of a voxel remains. Temporal units need sufficient numbers of observation to yield intervoxel variation. As we chose 1-minute as unit time, we observed 84 acquisitions (0.72 sec each) when using HCP data and 30 acquisitions (2 sec each) when using Kirby weekly data to calculate intervoxel correlation. 3- seconds shift in HCP or 2- seconds shift in Kirby weekly made 280 time-bins or 166 time-bins be observed for the time-varying changes, respectively. Though 1-minute bin per each database was obviously much smaller than 15 min (HCP) or 7 min (Kirby weekly), we still automatically assumed that 1- minute bin is stationary. Small time shifts allowed partially-overlapping observation of the time-varying changes. This strategy of defining voxels and time-bins as inputs revealed the slow temporal progresses of *k*_max_-core voxels-IC composition and interestingly the state transitions. In state transition, *k*_max_-core voxels-IC composition was abruptly changed so that the visual evaluation could count the numbers of state transitions. States were observed 2 to 21 or 4 to 20 times per study of HCP or Kirby weekly, respectively, whose counting was reproducible on visual reading (Fig. S4).

In 6x6x6 mm^3^, there was expected 20 million cells whose roles are either excitatory or inhibitory (*20*). Recently established distribution of 32 subtypes of excitatory/inhibitory subtypes of neurons revealed many overlaps but with definitive localization patterns (*26*). 20 million neurons in voxel units of this study has enough reason to act as voxel-nodes for positive correlation with other voxel-nodes and as voxel-nodes for negative correlation with other voxel- nodes. Once we found the symmetry of distribution of intervoxel correlation (Fig. S7), we wondered whether the anti-correlation (negative correlation) would reveal the undisclosed meaning of inhibitory network incorporated in the observed correlation matrix. In this preliminary endeavor, we simply used the absolute values of negative intervoxel correlation to compose another network, i.e. anti-correlation subnetwork, which were put into scale-free thresholding/*k*-core percolation similarly to correlation subnetwork. The output derived from this anticorrelation subnetwork were visualized and compared within individuals or within sessions in-an-individual with that from correlation network. Except a few exceptions, generally, *k*_max_- core voxels-IC composition of anticorrelation brain networks showed slow progress of *k*_max_-core voxels-IC composition of mostly distributed pattern and rare state shifts. In negative correlation analysis, larger number of participating voxels, evenly recruited voxels from almost all the ICs, were flickering in brain-rendered *k*_max_-core voxels along the 280 or 166 time-bins of individuals or sessions of an individual. Squeezing huge number of neurons and their mono- to poly-synaptic or electrical axonal transmission of activation (or information) makes intervoxel connectivity both positive and negative or cancelled (looking nil) making the functional connectivity matrix sparse. Whatever sparse intervoxel connectivity matrix look, considering the collective huge number of neurons in a voxel and thus all the voxels, the functional connectivity network is all-to-all linkage network, only flickering on the 1-minute time bins intermittently and alternatively. That is to say, voxels as spatial units of functional connectivity needs to be considered to have sustaining identities, impossible to be removed, merged or ignored ever. Incorporating the voxel-based negative functional correlation to positive one emphasized that we are observing many body all-to-all interacting system with N (number of voxels)-dimension. Spatiotemporal entity of N voxels observed by 1-minute time bins is the object of interest to be represented and visualized on animation to disclose their significance to represent time-varying progress of mental states of individuals and sessions of an individual.

The difference of the flickering pattern on brain-rendered images of positive and negative correlation study raised the possibility that both would be coordinated by different neuron subtypes, crudely excitatory and inhibitory. Inter-neuronal communication takes place electrically within neurons themselves having varying-length represented in their nomenclature of Exc_L5.6.IT (intra-telencephalic), Exc_L5.PT (pyramidal tract), or Exc_L6.CT (corticothalamic), and between neurons of glutamatergic or GABAergic synapses, which determines the speed of information transportation over the axons or across synapses. We cannot attend to sequential or orderly activation/inhibition of individual neurons within a voxel but observe the average 20 million neurons in a voxel during a 1-minute bin. Thus, we only observe the collective signal, and with blood oxygen level dependent (BOLD) signals of fMRI, we do not have chance to debate the theoretical issues of correlation-coordination frames of interpreting the intervoxel time-delay connections of the brain (*27*), which are found useful for interpreting electrical signals on electroencephalography or magnetoencephalography (*28*). Nervetheless, in our study, we found that spatiotemporal quanta of voxel/minute time-bin allowed the observation of time-varying changes of mental states at rest, which had been considered drifting, fluctuating, open (cannot be sequestered even in the non-activation resting state paradigm of fMRI acquisition) (*29*). We had the fortune to discover abrupt transition of states as would-be surrogates of mental states amid more prevalent slower temporal progresses of coreness *k*-values or *k*_max_-core voxels. In this study, we used the scheme to annotate the voxels, especially *k*_max_-core voxels to ICs derived from group ICA of 180 subjects of HCP database. Coloring and animation enabled the readers to read out the visual findings immediately, simply and easily. Counting and expressing the population prevalence of *k*_max_-core voxels-IC composition and transforming the individual’s data in animation format to intermittent snapshots was just the alternative way of presenting the findings.

*k*-core percolation enabled eloquent disclosure of hierarchical structure of functional intervoxel connectivity upon the image data of rsfMRI. In our previous study (*23*), using static study of hyperbolic embedding (and thus confirmation of scale-freeness of voxels’ degree distribution after thresholding) and associated *k*-core percolation, we attended most to the *k*_max_- core voxels across 30 individuals, and in this study, we extended the number of individuals to 180 inclusive of those 30. Easily extended to dynamic study, meaning that we observed 280 sequentially-followed observation with overlaps per individual, and thus around 50,000 bins of unit states were scrutinized visually. Totally 2,000 or more were labeled as states (1,500 for 25,000 in case of Kirby weekly), and thus 25 bins per state (16 bins, Kirby weekly). Belonging of *k*_max_-core voxels to ICs and thus composition could best be viewed on stacked histogram on animation and also brain-rendered images on animation. On the other hand, if one is interested in the position of every voxel along the hierarchical structure of the functional intervoxel connectivity especially time-varying on animation, flagplots on animation and voxel coreness *k*- values brain maps on animation are the best choice. Flagplots on animation of the data from HCP and Kirby weekly were investigated in detail here. Interestingly, flagplots on animation showed the concordant time-varying changes as raw plots of *k*-core percolation. The gradual lengthening and shortening of the horizontal axis of flagplots on animation was a good parameter of the deepening and shallowing of the hierarchical structures in toto, and the flagplots on animation revealed the separate behavior of voxels’ belonging to ICs in the link of their IC compositions.

Data were managed to be summed up as excel or mat files for possible further analysis. 5,937x280 (or x166) matrix per individual or per session represented the coreness *k*-values of the entire brain voxels and the signature of each individual (or that session of an individual). As any confirmatory comparison standard is currently unavailable, based on educated guess after seeing dozens of thousands bins or hundreds of outputs on animation from individuals/sessions we propose that voxels’ time-varying changes are related with the time-varying mental states and further their global intervoxel information processing. Obviously, we did observe intervoxel connectivity pairwise for _5,937_C_2_ combination and visualized on hyperbolic disc (*23, 24*), we took a step further to take advantage of simplicity of *k*-core percolation, and then taking further steps to reach finally voxels’ coreness *k*-values mapping (Fig S6). This was an important dimension reduction just as we did in our previous study (*30*) using directed networks of functional inter- nodal connectivity using rat PET data, i.e. afferent node capacity and efferent node capacity per nodes. Possibility of comparison between normal population and an individual of interest using any of the above parameters on animation or off animation (static) are to be studied further.

Pulses and *k*_max_-core voxels, found unexpectedly, raised much interest. Scrutinizing the characteristics of pulses showing up 6.4 times per 15 minutes in HCP data, and 4.8 times per 7 minutes in Kirby weekly data disclosed the findings that 1) pulses showed up anywhere within the states or just before or on the state transitions, mostly single but sometimes in train of singles, 2) frequently the pulses heralded the following state transition, but are not always followed by a state transition, and 3) pulses were composed of voxels belonging almost to all of the ICs (from DMN to UNC) which mimicked the distributed pattern on stacked histogram on animation. However, on flagplots on animation, pulses showed the short or shortened horizontal length, to let us infer that the pulse heralds the state transition, either successful or unsuccessful, mostly with decreased depth of global hierarchy as well as IC-wise hierarchy. Pulse can be considered as Dirac delta function on the temporal axis of voxels’ marching on, signifying the collective efforts of the brain to change the state. If this trial is successful, integral of delta function Heaviside function give birth to state transition. This explanation was just an analogy based on the presumed raison d’etre of pulses, intermittently observed on stacked histogram on animation in any individual of HCP data or in any session of an individual of Kirby weekly datasets.

Intriguing question roamed around regarding whether the instantaneous correlation of each time bin (1-minute) is either just the noisy fluctuation or functionally meaningful state representation (*7, 9, 31, 32*). Based on our observation in this study, on the tens of thousands of time-bins from normal adult humans, we do not believe that what we observed in our study could be made from noise (*14*), however, we are not going to insist that every combination of voxels- IC composition at every time bin means significant for behavior, either implicit or explicit. The heterogeneity of mental states could not be described by other methods such as introspective narratives, either self or from instrumental recordings. The correlative behavior and time-varying hierarchical constitution of core voxels remains to be understood (*8, 28*), considering that 7 to 15-minutes observation is just one session of the individual, that static is a longer-standing dynamic which did not seem to represent simple integration of the 1-minute bins in any easy way, and that another session in another time yielded the similar or the different uppermost voxels-IC composition pattern in an individual. Individual differences of coreness *k*-value map on animation or stacked histogram plots on animation resided both in the unique character of the subject individual and in the session of the study of an individual. Commonality can thus be defined as norms for control individuals, then we might be able to detect the anomaly.

Uniqueness of signature can be annotated for an individual on the analysis of just one session of each individual or of many repeated sequential sessions of that individual. With this information kept in mind, we might now dare to fingerprint of an individual or at least a specific session of a designated individual. Whether this objective is achieved by the methods of this study remains to be found.

Prominent differences of mental states can be achieved by anesthesia, sedation with drugs, sleep, and in the states of the altered consciousness in humans as well as in animals (*33–36*). Our method is expected to elucidate the rsfMRI surrogate results for time-varying mental states of the subject individual humans under anesthesia, sleep or in altered consciousness.

Temporal progress and state transition using spatiotemporal bins of voxel-minute are proposed to be applied for these studies. Correlation/anti-correlation does not yield directed functional connectivity graphs for functional connectivity. Though *k*-core percolation and ensuing plots/maps on animation visualization made the edge information of the brain network to the voxel-node representation, we still use the instantaneous collective correlation not allowing any periodicity/delay for interpretation of intervoxel relationship. And thus another strategy of understanding all-to-all voxels’ interaction including higher order rather than pairwise interaction and the concomitant coordination/propagation is still beyond the current study. However, at least, considering that the spatiotemporal bins are large enough to show the collective phenomena of so many neurons during such a long time 1-min from the intervoxel interaction, we rather drop the notion that sequential activation/propagation of spontaneously firing units at resting state, but adopt better idea that every voxel is an independent generator to produce autonomously only to communicate without any blocking (in our designated time-space frames) to yield collective multistable-metastable-synchronized-transient outputs at our read-outs, even coming to look like oscillating at the global/gross level (*17, 18, 37, 38*). We should not forget the role of anti-correlation, which was long ignored, and is derived from the inhibitory neurons mainly wired as interneurons, and we need to develop the synthesis method of the current study of separately observed positive and (absolute-valued) negative intervoxel correlations. One last issue is about the model itself, as investigators used real values for random matrix, what if the entries of matrix is complex values (u+vi) for non-Hermitian random matrix (*39, 40*), we might integrate the information in the progress of state and pulse along the time axis so that we integrate the phase (angle) for coherence and the amplitude (envelope) for correlation. In this investigation, we used voxels and time-bins, adopted the thresholding method originally developed for hyperbolic embedding, simple and easy but recursive filtration (percolation) of adjacency matrix and finally made the easy-to-read plots/maps both for coreness *k*-values and for *k*_max_-core voxels elucidating hierarchical structures of the functional resting intervoxel connectivity. Visualization using scale-free skeleton of correlation matrix, static for status and dynamic for time-varying on animation, including absolute negative correlations for anti-correlated subnetworks is the key advance of this investigation. Individual voxel-based interpretation is now available for clinical translation of resting-state fMRI for characterization of mental states of humans having brain disorders difficult to define and to be put into groups. Individual interpretation using parametric mapping/multiple comparison/disclosure of local connectivity anomaly with correlation/anti-correlation better with the aid of deep learning of variational autoencoder and the similar will be the next step and the limitation of the current status of this investigation.

## MATERIALS AND METHODS

### Data preprocessing

We downloaded rsfMRI data from the Human connectome project (www.humanconnectomeproject.org). We included 180 participants aged 22 to 36 without any significant history of psychiatric disorder, neurological or cardiovascular disease (*41, 42*). We used minimally preprocessed rsfMRI data (*43*) and performed further preprocessing: smoothing using 6 mm full-width at half maximum of Gaussian kernel, bandpass filtering (0.01 Hz – 0.1 Hz). Next, we downsampled the data from 91 x 109 x 91 dimension with 2 x 2 x 2 mm size- voxel to 31 x 37 x 31 dimension with 6 x 6 x 6 mm^3^ size-voxel, so that we can reduce computational load. We applied a mask to exclude voxels that do not belong to the brain, resulting in 5,937 voxels that are included in this analysis.

We downloaded rsfMRI data from the Kirby weekly (*44*). We included all the 156 sessions of an individual in this project and performed preprocessing. The motion parameters were estimated and despiking was performed using BrainWavelet Toolbox (*45*). The registration to the standard space was performed to MNI space after slice time correction and realignment.

The registered images were smoothed with 8 mm Gaussian kernel and regressed out of 6 motion parameters, white and CSF signals. The temporal filtering with 0.01 to 0.1 Hz and the down- sampling were just like those of HCP data to include 5,937 voxels with 6 x 6 x 6 mm^3^.

### Independent component analysis

We carried out independent component analysis (ICA) to find ICs, i.e., the resting-state networks, using Multivariate Exploratory Linear Decomposition into Independent Components (MELODIC) (*46*). We obtained spatial maps of ICs, and applied threshold (Z > 6) to generate binary masks. In this study, we included 15 ICs: aDMN (anterior default mode network), DMN, PCN (precuneus network, equivalent to posterior DMN), SN (salience network), DAN (dorsal attention network), L CEN (left central executive network), R CEN (right CEN), SMN (sensorimotor network) 1, SMN2, AN (auditory network), VN (visual network) 1, VN2, VN3, VN4, and VAN (ventral attention network) (Fig. S1). We classified them into seven categories for comprehensibility: DMN, SN, DAN, CEN, SMN, AN, and VN and the remainders UNC.

### Dynamic data analysis: sliding-window analysis

For HCP data, we used sliding-window analysis to investigate the nonstationary and time-dependent dynamics of the brain connectivity. The window size was set close to 1 minute (84 volumes, 60.48 sec) with a shift of 4 volumes (2.88 sec), resulting in 280 windows. A connectivity matrix of each window was calculated to conduct *k*-core percolation. We implemented *k*-core percolation for each connectivity matrix after applying the threshold, which meets the scale-freeness of the network, to generate a binary matrix. Edges with values greater than 0.65 were assigned as one and otherwise 0.

For Kirby weekly data, window size was set to 1 minute (30 volumes) with a shift of 2 sec. A connectivity matrix of each window was calculated using instantaneous correlation. Edges with values greater than 0.8 were assigned as one and otherwise 0 to make adjacency matrix.

For individuals and sessions of both studies, the same threshold value per study was used for individuals and for sessions, respectively, with the criteria we reported previously (*23*) to procure as many voxels while guaranteeing scale-freeness. There were range of thresholds meeting this criteria among which we chose ones (0.65 for HCP and 0.80 for Kirby weekly), and with these set thresholds, nodes included more than 72% of the voxels (mean 88.5% of 5,937 total voxels) for HCP bins (180 individuals x 280 bins).

### k-core percolation

We conducted *k*-core percolation to investigate an individual’s hierarchical core structure of a functional brain network (*23, 25*). Metaphorically, this method peels the layer of the network as if it were an onion. This procedure first removes nodes of degree 1 (*k* = 1). As nodes are removed, the degree of the remaining nodes also changes. Nodes, whose degrees were higher, are eventually removed if they meet the removal criteria while *k* increases. The procedure is performed recursively by incrementing *k* by 1 until no further process is possible. A subgraph, called *k*-core, is obtained when all nodes with degree less than *k* is removed. The last surviving core was called *k*_max_-core. After *k*-core percolation was performed on each subject’s data, we classified *k*_max_-core voxels using IC maps. All the voxels were given the *k* values of the step when they were removed. This was called coreness *k* value of each voxel.

For dynamic study, 280 time-bins of HCP individuals and 156 time-bins of Kirby weekly sessions were put in to the *k*-core percolation. The coreness *k* values were used for making flagplots and brain-rendered coreness *k* images on animation and *k*_max_-core voxels-ICs compositions for making stacked histogram and brain-rendered *k*_max_-core images on animation (Fig. 1)

## Acknowledgments

We thank Drs. Seunggyun Ha and Chun-Kee Chung for helpful discussion.

## Author contributions

Conceptualization: DSL, HK

Methodology: YH, YKK, WW, HK, DSL

Investigation: YH, YKK, WW, HK, DSL

Visualization: YH, YKK, WW, HK, DSL

Funding acquisition: HK, DSL

Project administration: HK, DSL

Supervision: HK, DSL

Writing – original draft: YH, DSL

Writing – review & editing: HK, DSL

## Data and materials availability

All data are available in the main text or the supplementary materials. All data, code, and materials used in the analysis are available through a standard material transfer agreement with Seoul National University to academic and nonprofit by contacting the authors.

## List of Supplementary Materials

### Supplementary Figures

Fig. S1. Independent components produced using group ICA. Default mode network (DMN) in cyan, salience network (SN) in red, dorsal attention network (DAN) in orange, central executive network (CEN) in yellow, sensorimotor network (SMN) in green, auditory network (AN) in blue, visual network (VN) in violet and unclassified (UNC) in gray colors. Unclassified were the voxels remaining the ones designated, which included subcortex, cerebellum and others.

Fig. S2. Static analysis yielded 5,937x1 vector data which were put in to make flagplots and brain-rendered coreness *k-*values brain map. Dynamic analysis yielded 5,937x280 matrix of coreness *k-*values, which were transformed to make flagplots on animation and coreness *k*-values brain maps on animation.

Fig. S3. In this individual, brain-rendered images of positive connectivity showed asymmetry of *k*_max_-core voxels of DMN, DAN, CEN in frontal areas on animation. Only the state of DMN/CEN dominant, but not the VN dominant in positive connectivity showed asymmetry and not in negative connectivity.

Fig. S4. Reproducibility of visual counting of state transitions on the data of positive connectivity of HCP (n=180) individuals and of Kirby weekly (n=156) sessions in an individual.

Fig. S5. In a representative individual of HCP database, *k-*core percolation results of 280 time-bins were shown. This shows downward changes of coreness *k-*values of entire voxels according to steps of the percolation, gradual in general but with intermittent abrupt (explosive) decrease of voxels, which finally reached the uppermost level representing the *k*_max_-core value. The number of voxels remaining at the last step of *k-*core percolation ranged from several tens to almost a thousand.

Fig. S6. Coreness *k* values of the entire voxels were displayed on the transaxial templates of 10 slices of the MRI templates. As voxels’ coreness *k* values are the characteristic numbers of the voxels after due consideration of edge information (correlation or anti-correlation), pairwise edge information was translated successfully into node-voxel information. As they look like topography, voxel coreness *k* values can now be used for further analysis of comparison of an individual or a session with those of preset norms of individuals or preset norms of sessions.

Fig. S7. Correlation histogram of an individual from HCP project. A) Using initial voxel size of 2x2x2 mm^3^, -174,610 voxels could be used as input to yield weights of 15 giga undirected edges as correlation/ anti-correlation. 2) Using 6x6x6 mm^3^, 5,937 voxels as inputs to 17.6 mega edges. 3) Using 274 volumes of interest (VOIs), 274 voxels as 37 kilo edges.

### Supplementary Movies

Movie S1 . Brain-rendered image of *k*_max_-core voxels, stacked histogram, bar plot and flagplots in an individual of HCP on static data analysis. Brain rendered images, stacked histogram, bar plots and flagplots on animation on dynamic data analysis.

Movie S2. Brain-rendered image of *k*_max_-core voxels, stacked histogram, bar plot and flagplots of a session from an individual of Kirby weekly on static data analysis. Brain rendered images, stacked histogram, bar plots and flagplots on animation on dynamic data analysis.

Movie S3. Voxel coreness *k* values brain-rendered map and flagplots of two cases of HCP. Upper row of brain-rendered images on animation using 10 among 31 slices and middle row left flagplots on animation showed time-varying coreness *k* values. On stacked histogram of k-_max_ core voxels-IC compostion of this individual, combinations of dominant patterns were observed time-varyingly. Lower row of brain-rendered images and middle row right flagplots on animation showed different pattern in another individual, whose stacked histogram on animation showed distributed/DMN-CEN co-dominant pattern.

Movie S4. In an individual, positive connectivity on the left, negative connectivity on the right. Positive connectivity showed DMN/CEN dominant, visual dominant and distributed patterns ran in train, taking turns with quick state transition, interspersed with each other. Slow progress of *k*_max_-core voxels-ICs composition were observed and 16 state transitions showed up. Negative connectivity showed mostly distributed pattern with intermittent pulses of more complete composition. Positive and negative connectivity were different in the *k*_max_-core voxels-ICs composition.

Movie S5. In another individual, the different pattern of *k*_max_-core voxels-ICs composition between positive and negative connectivity was also observed. Again 15 state transitions were found with many pulses successful or unsuccessful for state transition. Interestingly, those pulses of positive and negative connectivity were different on brain-rendered images. *k*_max_-core voxels of the negative connectivity were with more widely/evenly distributed composed by larger numbers of voxels than those of positive connectivity.

Movie S6. Brain rendered images of *k*_max_-core voxels and stacked histogram on animation. Left animated images were from the analysis using positive correlation. Voxels belonging to DMN, DAN and CEN of both frontal areas showed transients of asymmetry of *k*_max_-core features on animation. Animated images on the right side were from negative correlation. No asymmetry was observed in the pattern of flickering voxels on animation.

Movie S7. Flagplot showing time-varying changes of hierarchical height for negative correlation. The length of *k*-core percolation changed over time independently from the total number of *k*_max_- core voxels. In both individuals, negative correlation showed distributed pattern of *k*_max_-core voxels-ICs composition, which showed plain progress with intermittent pulses/trains with larger number of *k*_max_-core voxels. Flags’ lengths on flagplots representing hierarchical height/depth waxed and waned repeatedly.

Movie S8. Comparison of static and dynamic studies. In this individual, in static study using positive intervoxel correlation, the pattern of DMN/CEN dominance was found in *k*_max_-core composition, however, in dynamic study other patterns including VN dominant, distributed and their combinations showed up with quick switches between themselves. State progress and abrupt transition made the vertical switching pattern, which was named the state transition. Distributed pattern of static study using negative correlation, looked proportional to the occupancy of IC-voxels and their sum, tentatively called horizontal summing-up pattern.

Movie S9. In another individual, the VN dominance pattern of the static study of positive correlation, though on dynamic study the half of the period was occupied by VN dominance pattern in *k*_max_-core composition consisting of five short fragments interspersed with the same amount of DMN/CEN co-dominance pattern. Distributed pattern of static study using negative correlation revealed similar phenominon to that of the case of Movie S8.

Movie S10. In the other individual showing distributed pattern of the static study of positive correlation, distributed pattern (vertically segmented along the elapsing time) was dominant in dynamic study but also with a few scattered VN dominant or DMN/CEN dominant segments.

Movie S11. Four sessions of an individual (Kirby weekly) were analyzed for hierarchical structure of voxels-IC composition on *k*-core percolation using functional intervoxel positive and negative correlations. A session of DMN/CEN/DAN co-dominance pattern (positive) was with distributed pattern (negative). Another DMN/CEN with distributed pattern (positive) and two other VN plus pattern with distributed pattern (negative).

Movie S12. Another four sessions of an individual (Kirby weekly) were presented. A session of distributed pattern (positive) was with distributed pattern (negative) and another session of SN/SMN/AN (positive) was with distributed minus pattern (negative). Another session could be a combination of strips of various dominance but uniquely included UNC (including voxels of cerebellum) on positive correlation analysis. In both positive and negative correlation analyses, it is noted that UNC voxels including cerebellum were among the uppermost hierarchy on *k*-core percolation. The last one was DAN/CEN (positive)-distributed (negative) pattern pair.

