## supplementary figures and movies for "Time-varying hierarchical core voxels disclosed by *k*-core percolation on dynamic inter-voxel connectivity resting-state fMRI"

Fig. S1. Independent components produced using group ICA.

Fig. S2. Static analysis yielded 5,937x1 vector data which were put in to make flagplots and brain-rendered coreness *k*-values brain map.

Fig. S3. In this individual, brain-rendered images of positive connectivity showed asymmetry of $k_{max}$-core voxels of DMN, DAN, CEN in frontal areas on animation.

Fig. S4. Reproducibility of visual counting of phase transitions on the data of positive connectivity of HCP (n=180) individuals and of Kirby weekly (n=156) sessions in an individual.

Fig. S5. In a representative individual of HCP database, *k*-core percolation results of 280 time-bins were shown.

Fig. S6. Coreness *k* values of the entire voxels were displayed on the transaxial templates of 10 slices of the MRI templates.

Fig. S7. Correlation histogram of an individual from HCP project.

Figure S1

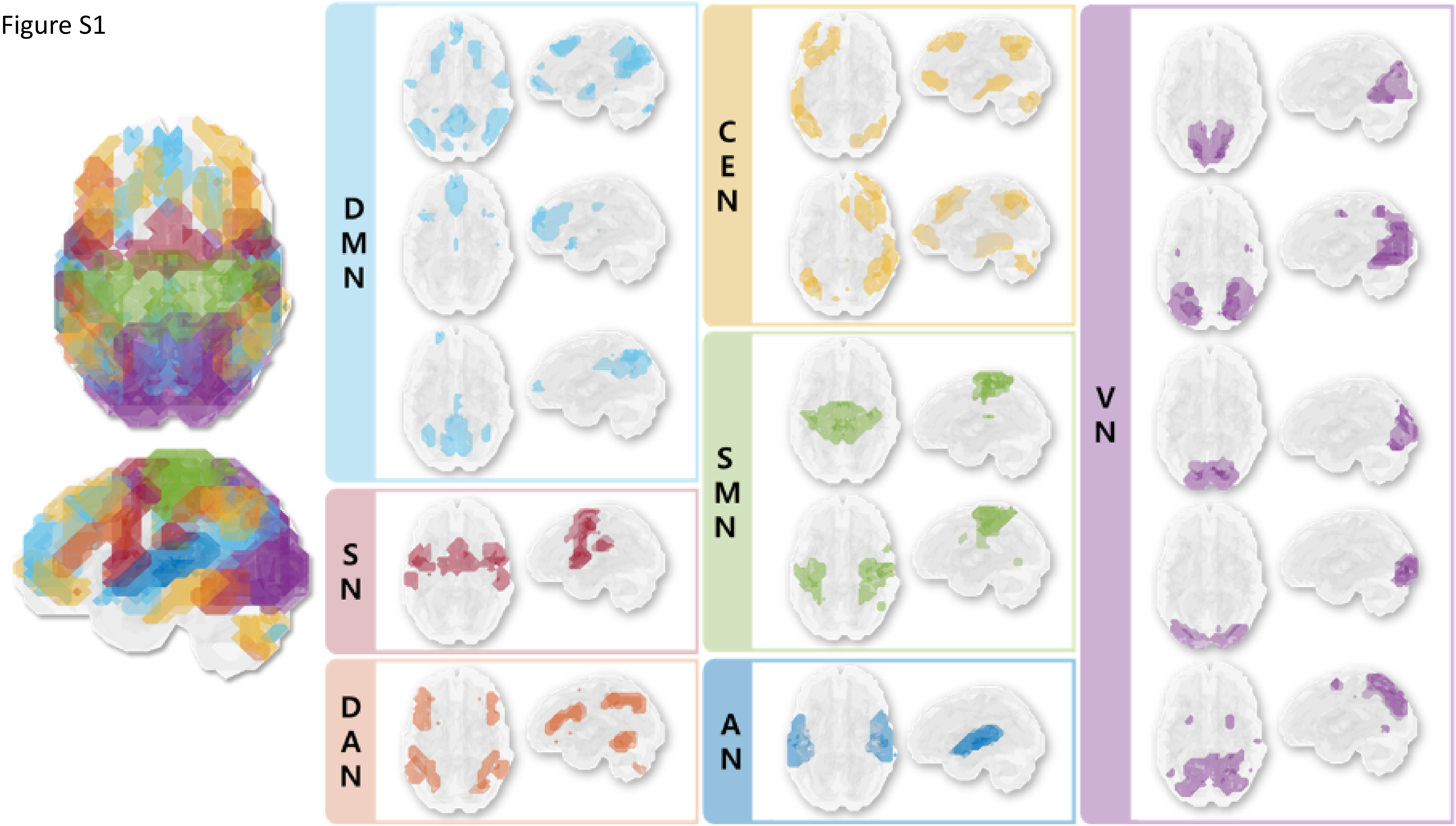

### Figure S2

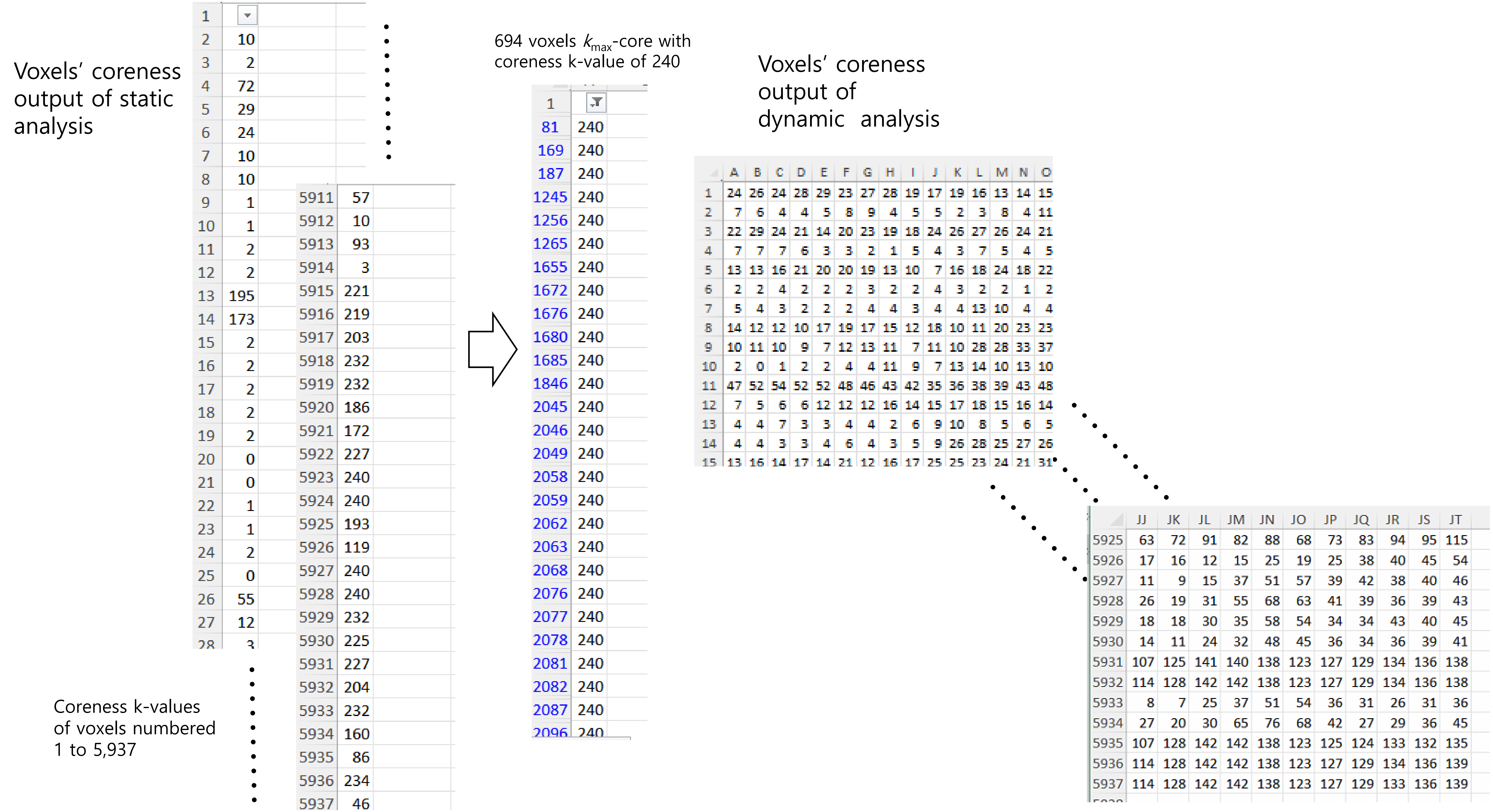


| | |
|---|---|
| 1 | |
| 2 | 10 |
| 3 | 2 |
| 4 | 72 |
| 5 | 29 |
| 6 | 24 |
| 7 | 10 |
| 8 | 10 |
| 9 | 1 |
| 10 | 1 |
| 11 | 2 |
| 12 | 2 |
| 13 | 195 |
| 14 | 173 |
| 15 | 2 |
| 16 | 2 |
| 17 | 2 |
| 18 | 2 |
| 19 | 2 |
| 20 | 0 |
| 21 | 0 |
| 22 | 1 |
| 23 | 1 |
| 24 | 2 |
| 25 | 0 |
| 26 | 55 |
| 27 | 12 |
| 28 | 3 |

| | |
|---|---|
| 5911 | 57 |
| 5912 | 10 |
| 5913 | 93 |
| 5914 | 3 |
| 5915 | 221 |
| 5916 | 219 |
| 5917 | 203 |
| 5918 | 232 |
| 5919 | 232 |
| 5920 | 186 |
| 5921 | 172 |
| 5922 | 227 |
| 5923 | 240 |
| 5924 | 240 |
| 5925 | 193 |
| 5926 | 119 |
| 5927 | 240 |
| 5928 | 240 |
| 5929 | 232 |
| 5930 | 225 |
| 5931 | 227 |
| 5932 | 204 |
| 5933 | 232 |
| 5934 | 160 |
| 5935 | 86 |
| 5936 | 234 |
| 5937 | 46 |

| | |
|---|---|
| 1 | |
| 81 | 240 |
| 169 | 240 |
| 187 | 240 |
| 1245 | 240 |
| 1256 | 240 |
| 1265 | 240 |
| 1655 | 240 |
| 1672 | 240 |
| 1676 | 240 |
| 1680 | 240 |
| 1685 | 240 |
| 1846 | 240 |
| 2045 | 240 |
| 2046 | 240 |
| 2049 | 240 |
| 2058 | 240 |
| 2059 | 240 |
| 2062 | 240 |
| 2063 | 240 |
| 2068 | 240 |
| 2076 | 240 |
| 2077 | 240 |
| 2078 | 240 |
| 2081 | 240 |
| 2082 | 240 |
| 2087 | 240 |
| 2096 | 240 |

| | A | B | C | D | E | F | G | H | I | J | K | L | M | N | O |
|---|---|---|---|---|---|---|---|---|---|---|---|---|---|---|---|
| 1 | 24 | 26 | 24 | 28 | 29 | 23 | 27 | 28 | 19 | 17 | 19 | 16 | 13 | 14 | 15 |
| 2 | 7 | 6 | 4 | 4 | 5 | 8 | 9 | 4 | 5 | 5 | 2 | 3 | 8 | 4 | 11 |
| 3 | 22 | 29 | 24 | 21 | 14 | 20 | 23 | 19 | 18 | 24 | 26 | 27 | 26 | 24 | 21 |
| 4 | 7 | 7 | 7 | 6 | 3 | 3 | 2 | 1 | 5 | 4 | 3 | 7 | 5 | 4 | 5 |
| 5 | 13 | 13 | 16 | 21 | 20 | 20 | 19 | 13 | 10 | 7 | 16 | 18 | 24 | 18 | 22 |
| 6 | 2 | 2 | 4 | 2 | 2 | 2 | 3 | 2 | 2 | 4 | 3 | 2 | 2 | 1 | 2 |
| 7 | 5 | 4 | 3 | 2 | 2 | 2 | 4 | 4 | 3 | 4 | 4 | 13 | 10 | 4 | 4 |
| 8 | 14 | 12 | 12 | 10 | 17 | 19 | 17 | 15 | 12 | 18 | 10 | 11 | 20 | 23 | 23 |
| 9 | 10 | 11 | 10 | 9 | 7 | 12 | 13 | 11 | 7 | 11 | 10 | 28 | 28 | 33 | 37 |
| 10 | 2 | 0 | 1 | 2 | 2 | 4 | 4 | 11 | 9 | 7 | 13 | 14 | 10 | 13 | 10 |
| 11 | 47 | 52 | 54 | 52 | 52 | 48 | 46 | 43 | 42 | 35 | 36 | 38 | 39 | 43 | 48 |
| 12 | 7 | 5 | 6 | 6 | 12 | 12 | 12 | 16 | 14 | 15 | 17 | 18 | 15 | 16 | 14 |
| 13 | 4 | 4 | 7 | 3 | 3 | 4 | 4 | 2 | 6 | 9 | 10 | 8 | 5 | 6 | 5 |
| 14 | 4 | 4 | 3 | 3 | 4 | 6 | 4 | 3 | 5 | 9 | 26 | 28 | 25 | 27 | 26 |
| 15 | 13 | 16 | 14 | 17 | 14 | 21 | 12 | 16 | 17 | 25 | 25 | 23 | 24 | 21 | 31 |

| | JJ | JK | JL | JM | JN | JO | JP | JQ | JR | JS | JT |
|---|---|---|---|---|---|---|---|---|---|---|---|
| 5925 | 63 | 72 | 91 | 82 | 88 | 68 | 73 | 83 | 94 | 95 | 115 |
| 5926 | 17 | 16 | 12 | 15 | 25 | 19 | 25 | 38 | 40 | 45 | 54 |
| 5927 | 11 | 9 | 15 | 37 | 51 | 57 | 39 | 42 | 38 | 40 | 46 |
| 5928 | 26 | 19 | 31 | 55 | 68 | 63 | 41 | 39 | 36 | 39 | 43 |
| 5929 | 18 | 18 | 30 | 35 | 58 | 54 | 34 | 34 | 43 | 40 | 45 |
| 5930 | 14 | 11 | 24 | 32 | 48 | 45 | 36 | 34 | 36 | 39 | 41 |
| 5931 | 107 | 125 | 141 | 140 | 138 | 123 | 127 | 129 | 134 | 136 | 138 |
| 5932 | 114 | 128 | 142 | 142 | 138 | 123 | 127 | 129 | 134 | 136 | 138 |
| 5933 | 8 | 7 | 25 | 37 | 51 | 54 | 36 | 31 | 26 | 31 | 36 |
| 5934 | 27 | 20 | 30 | 65 | 76 | 68 | 42 | 27 | 29 | 36 | 45 |
| 5935 | 107 | 128 | 142 | 142 | 138 | 123 | 125 | 124 | 133 | 132 | 135 |
| 5936 | 114 | 128 | 142 | 142 | 138 | 123 | 127 | 129 | 134 | 136 | 139 |
| 5937 | 114 | 128 | 142 | 142 | 138 | 123 | 127 | 129 | 133 | 136 | 139 |

Figure S3

### Asymmtery found on brain-rendered images

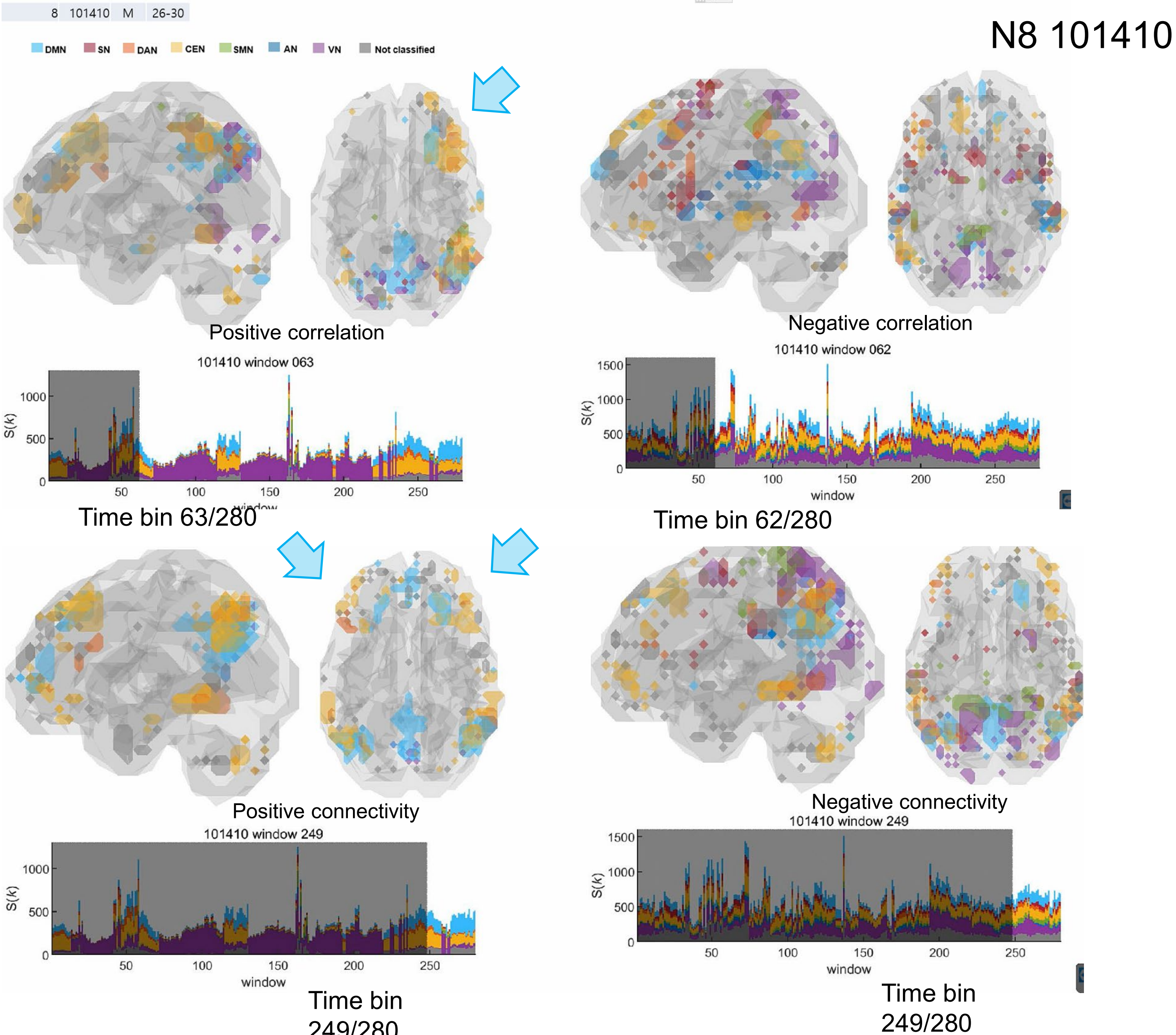

Figure S4

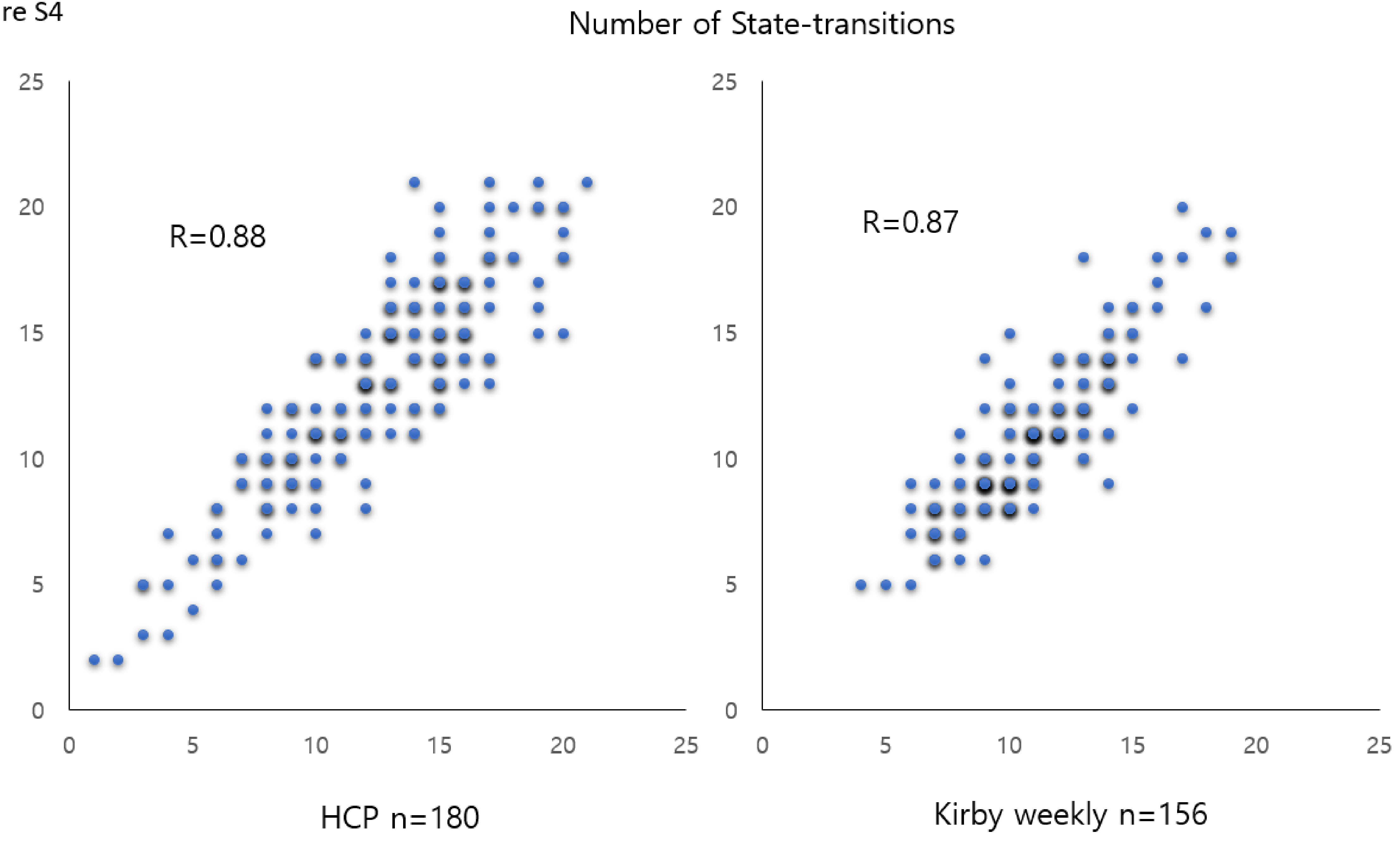
Number of State-transitions
R=0.88
HCP n=180
R=0.87
Kirby weekly n=156
0
5
10
15
20
25

Figure S5

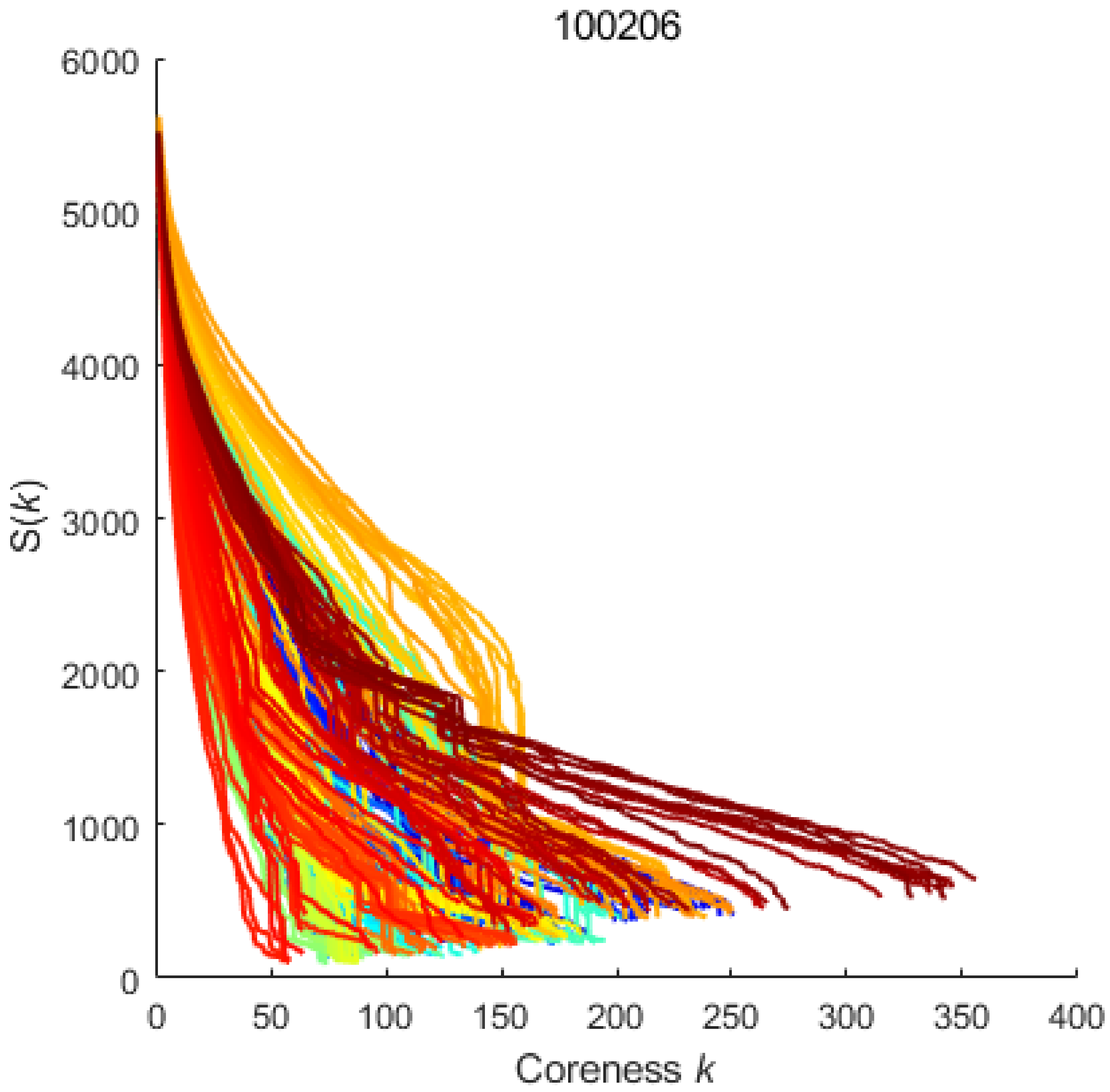

100206
6000
5000
4000
3000
2000
1000
0
S(k)
0
50
100
150
200
250
300
350
400
Coreness k

Figure S6

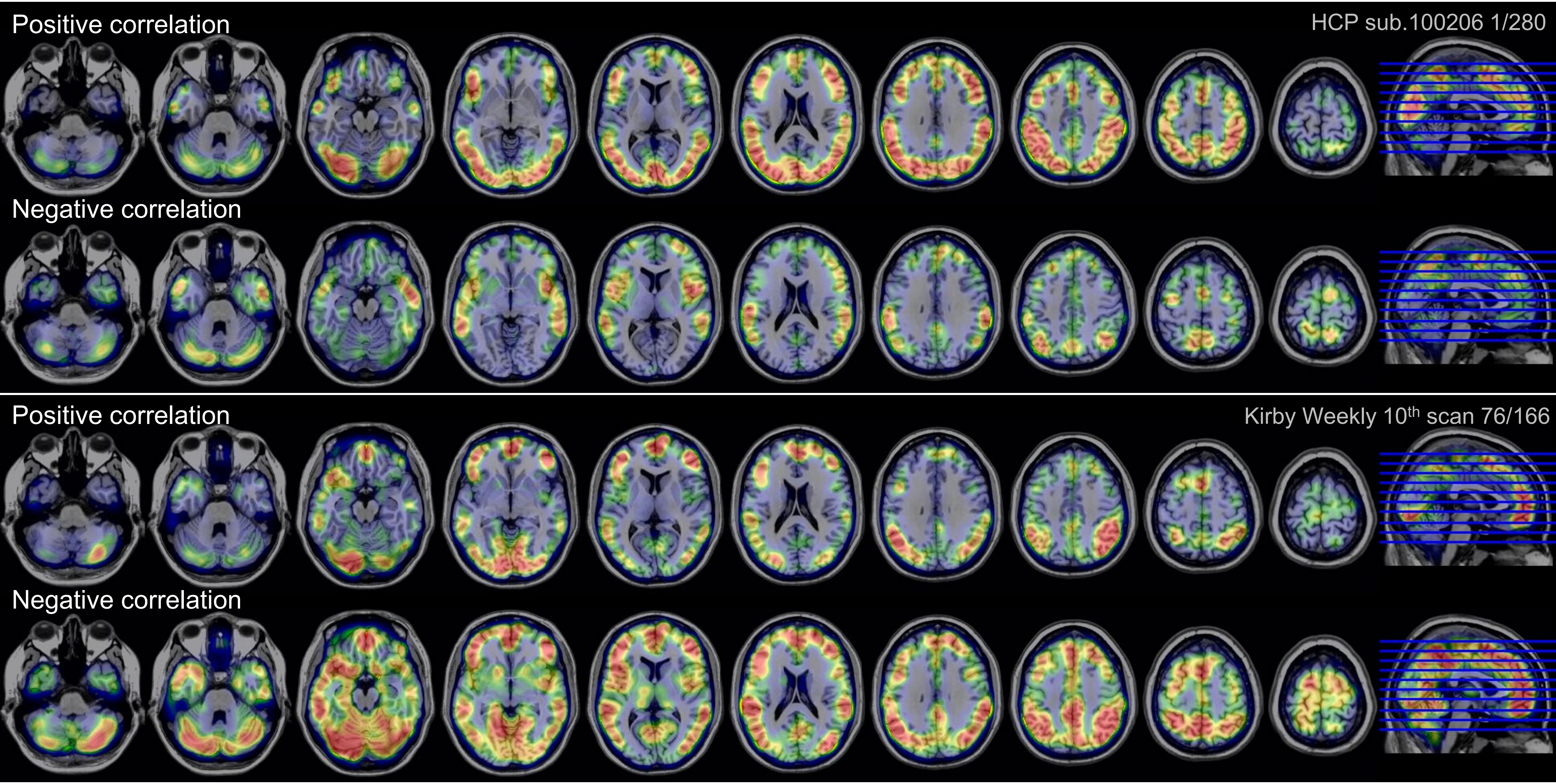
Positive correlation
HCP sub.100206 1/280
Negative correlation
Positive correlation
Kirby Weekly 10th scan 76/166
Negative correlation

### Figure S7

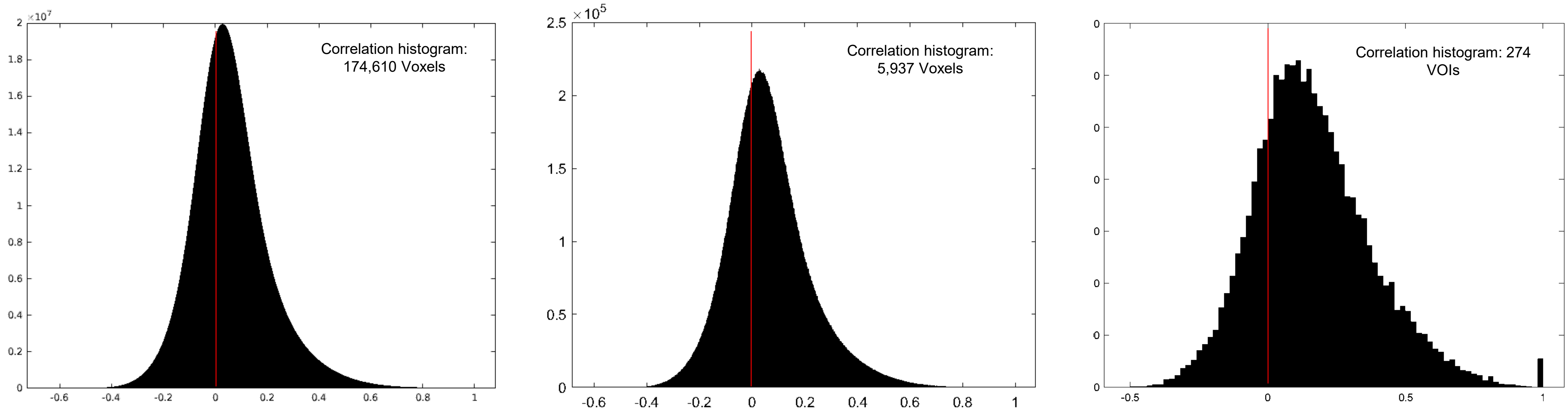

Correlation histogram: 174,610 Voxels
Correlation histogram: 5,937 Voxels
Correlation histogram: 274 VOIs

Supplementary Movies

**Title:** Time-varying hierarchical core voxels disclosed by k-core percolation on dynamic inter-voxel connectivity resting-state fMRI

**Authors:** Youngmin Huh[1], Yeon Koo Kang[2,3], Wonseok Whi[1,2,3,4], Hyekyoung Lee[5], Hyejin Kang[5]*, Dong Soo Lee[1,2,3]*

**Movie S1 . Brain-rendered image of $k_{max}$-core voxels, stacked histogram, bar plot and flag plots in an individual of HCP on static data analysis**. Brain rendered images, stacked histogram, bar plots and flag plots on animation on dynamic data analysis.

**Movie S2**. **Brain-rendered image of $k_{max}$-core voxels, stacked histogram, bar plot and flag plots of a session from an individual of Kirby weekly on static data analysis**. Brain rendered images, stacked histogram, bar plots and flag plots on animation on dynamic data analysis.

**Movie S3. Voxel coreness *k* values brain-rendered map and flagplots of two cases of HCP**. Upper row of brain-rendered images on animation using 11 among 31 slices and middle row left flagplots on animation showed time-varying coreness k values. On stacked histogram of k-$_{max}$ core voxels-IC compostion of this individual, combinations of dominant pattern was observed. Lower row of brain-rendered images and middle row right flagplots on animation showed another pattern, whose stacked histogram on animation showed distributed/DMN-CEN co-dominant pattern.

**Movie S8. Comparison of static and dynamic studies. In this individual, in static study using positive inter-voxel correlation, the pattern of DMN/CEN dominance was found in $k_{max}$-core composition, however, in dynamic study other patterns including VN dominant, distributed and their combinations showed up with quick switches. State progress and abrupt transition made the vertical switching pattern, which was also called state transition**. Distributed pattern of static study on the analysis using negative correlation, looked proportional to the occupancy of IC-voxels and their sum, tentatively called horizontal summing-up pattern.

**Movie S9. In another individual, the VN dominance pattern of the static study of positive correlation, though the half of the period was occupied by VN dominance pattern in $k_{max}$-core composition consisting of five short fragments interspersed with the same amount of DMN/CEN co-dominance pattern**. Distributed pattern of static study using negative correlation revealed similar situation to that of the case of Movie S7.

**Movie S10. In the other individual showing distributed pattern of the static study of positive correlation, distributed pattern (vertically segmented along the elapsing time) was dominant in dynamic study but also with a few scattered VN dominant or DMN/CEN dominant segments.**

**Movie S11. Four sessions of an individual (Kirby weekly) were analyzed for hierarchical structure of voxels-IC composition on *k*-core percolation using functional intervoxel positive and negative correlations**. A session of DMN/CEN/DAN co-dominance pattern (positive) was with distributed pattern (negative). Another DMN/CEN with distributed pattern (positive) and two other VN plus pattern with distributed pattern (negative).

**Movie S12. Another four sessions of an individual (Kirby weekly) were presented. A session of distributed pattern (positive) was with distributed pattern (negative) and another session of SN/SMN/AN (positive) was with distributed minus pattern (negative).** Another session could be a combination of strips of various dominance but uniquely included UNC (including voxels of cerebellum) on positive correlation analysis. In both positive and negative correlation analyses, it is noted that UNC voxels including cerebellum were among the uppermost hierarchy on *k*-core percolation. The last one was DAN/CEN (positive)-distributed (negative) pattern pair.

### Movie S1

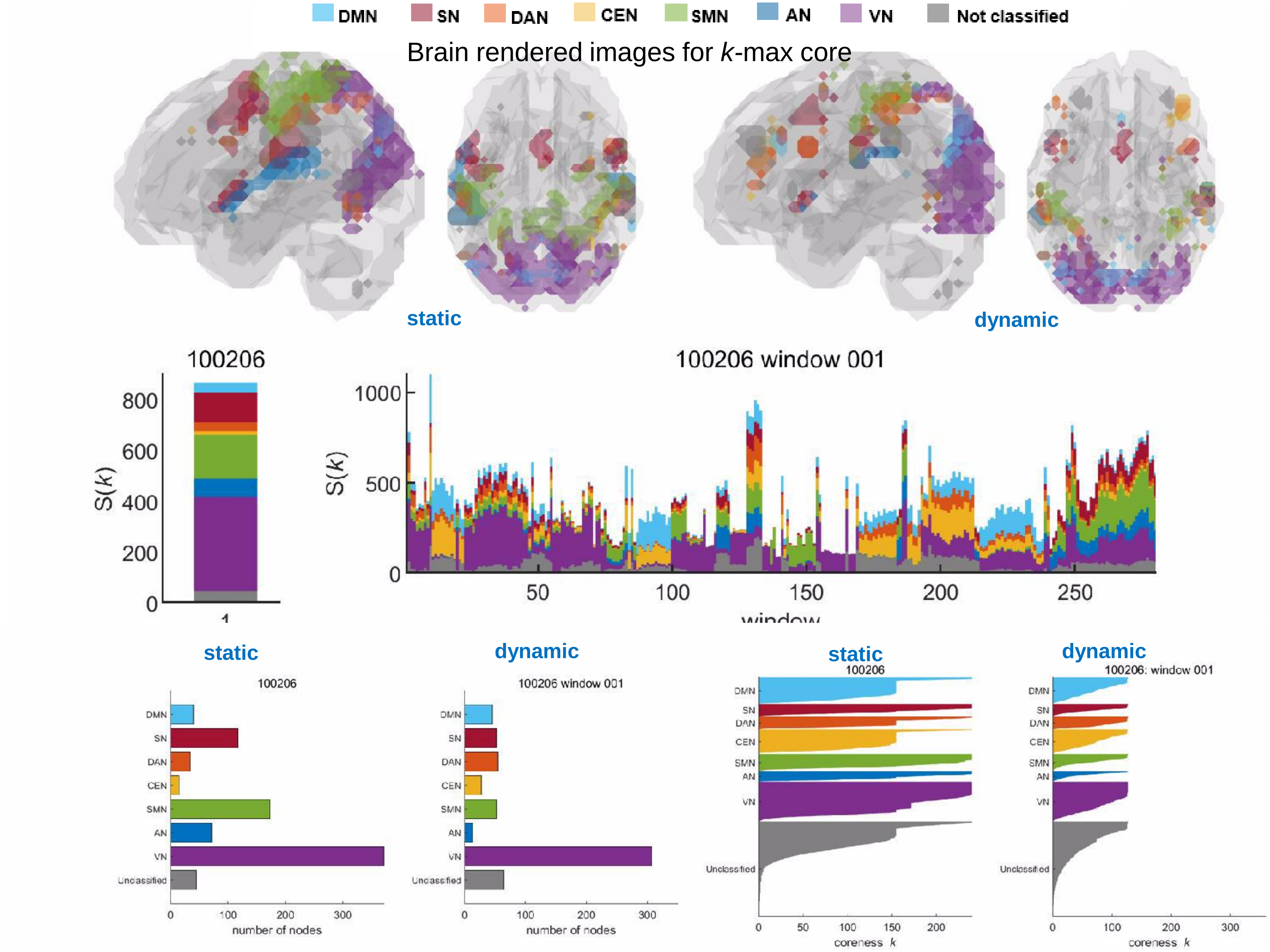


Barplot for *k*-max core

Flagplots for voxel coreness *k* values

Movie S2

2nd scan/ 156 weekly in a single subject (Kirby weekly)

DMN SN DAN CEN SMN AN VN Not classified

Brain rendered images for *k*-max core

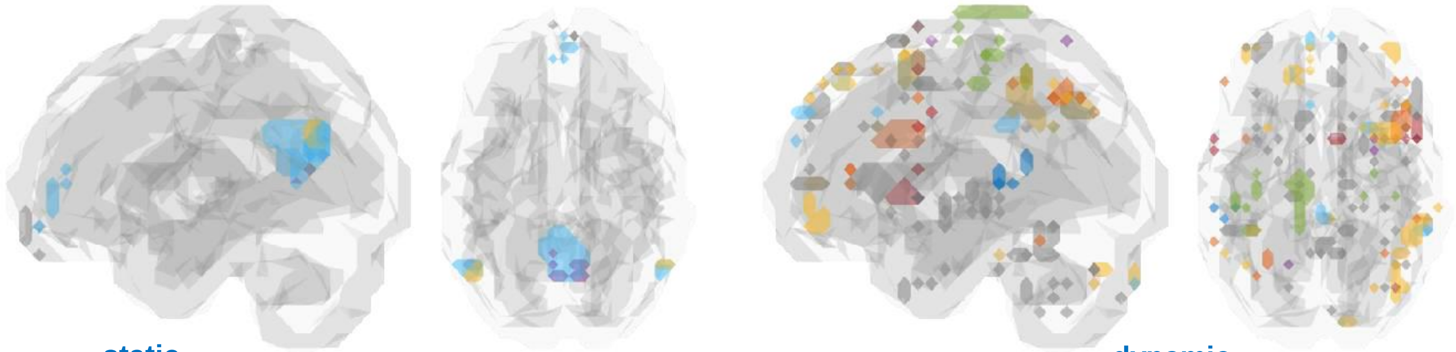

static dynamic

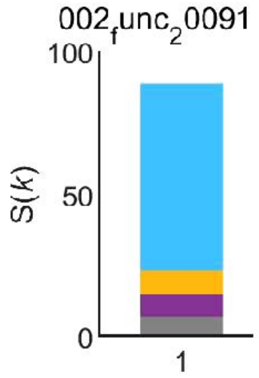


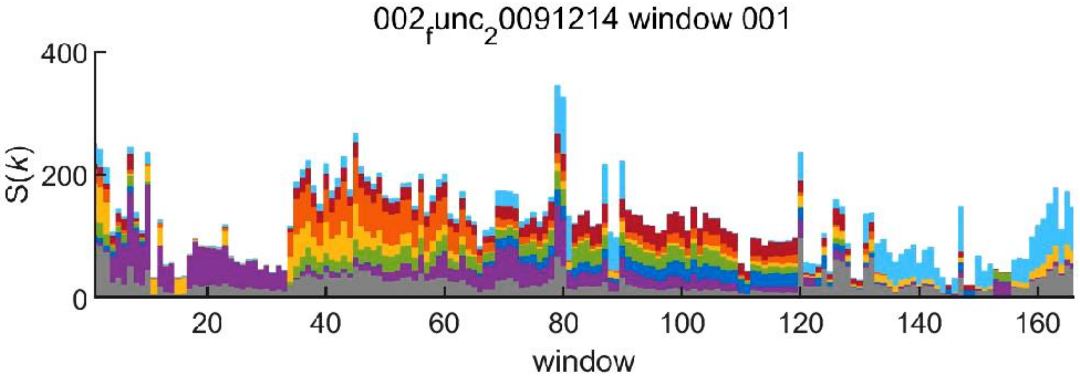


static dynamic static dynamic

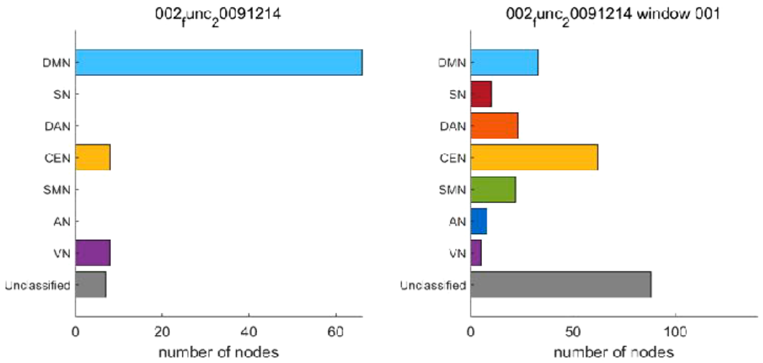


Barplot for *k*-max core

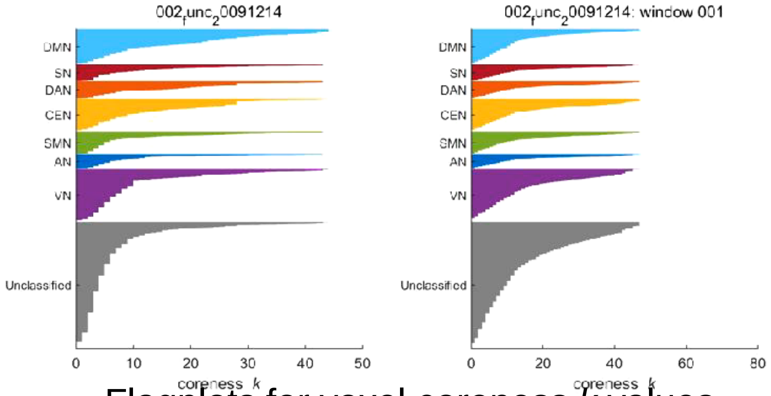


Flagplots for voxel coreness *k* values

Movie S3

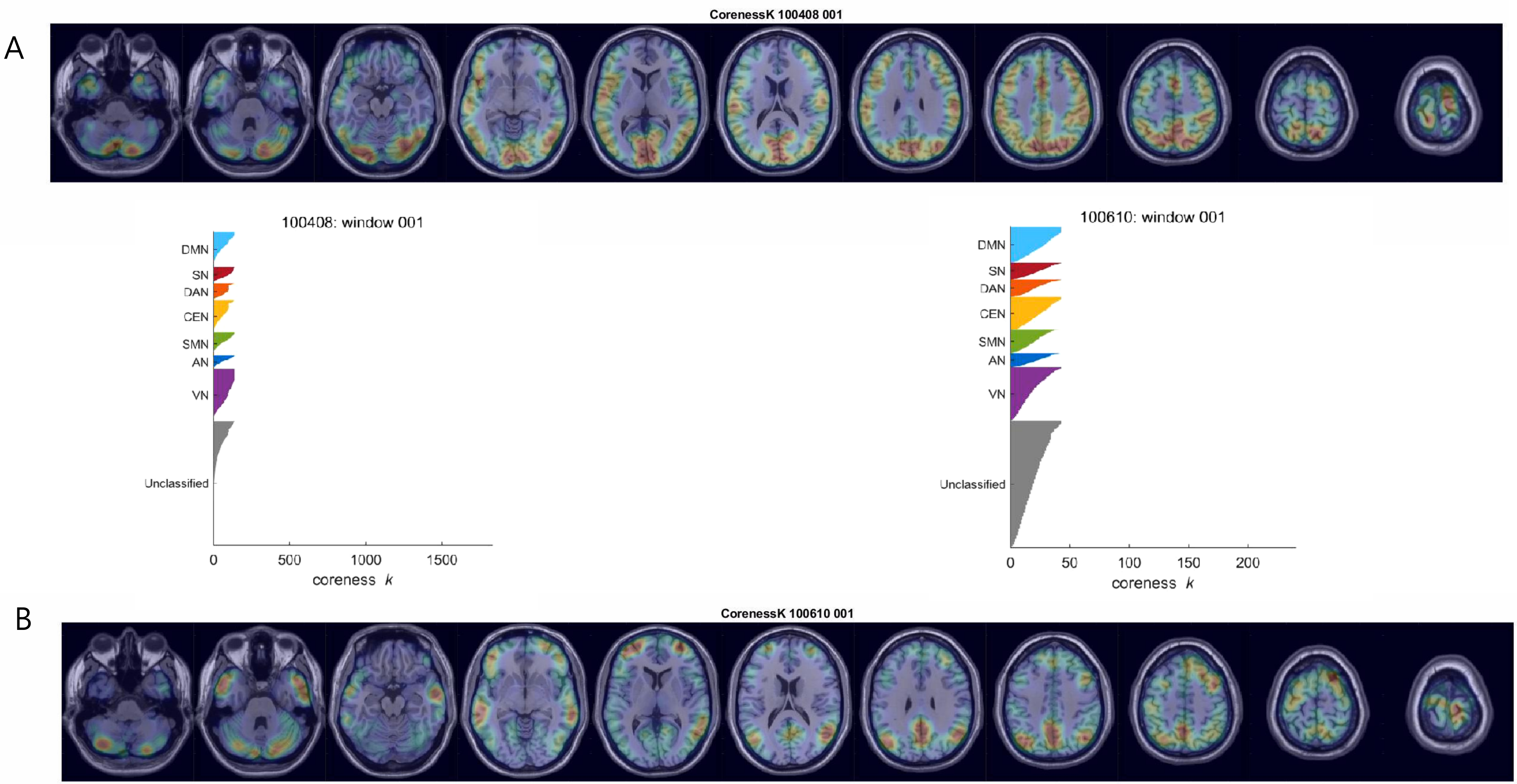

A
CorenessK 100408 001
100408: window 001
DMN
SN
DAN
CEN
SMN
AN
VN
Unclassified
0
500
1000
1500
coreness k
100610: window 001
DMN
SN
DAN
CEN
SMN
AN
VN
Unclassified
0
50
100
150
200
coreness k
B
CorenessK 100610 001

### Movie S4

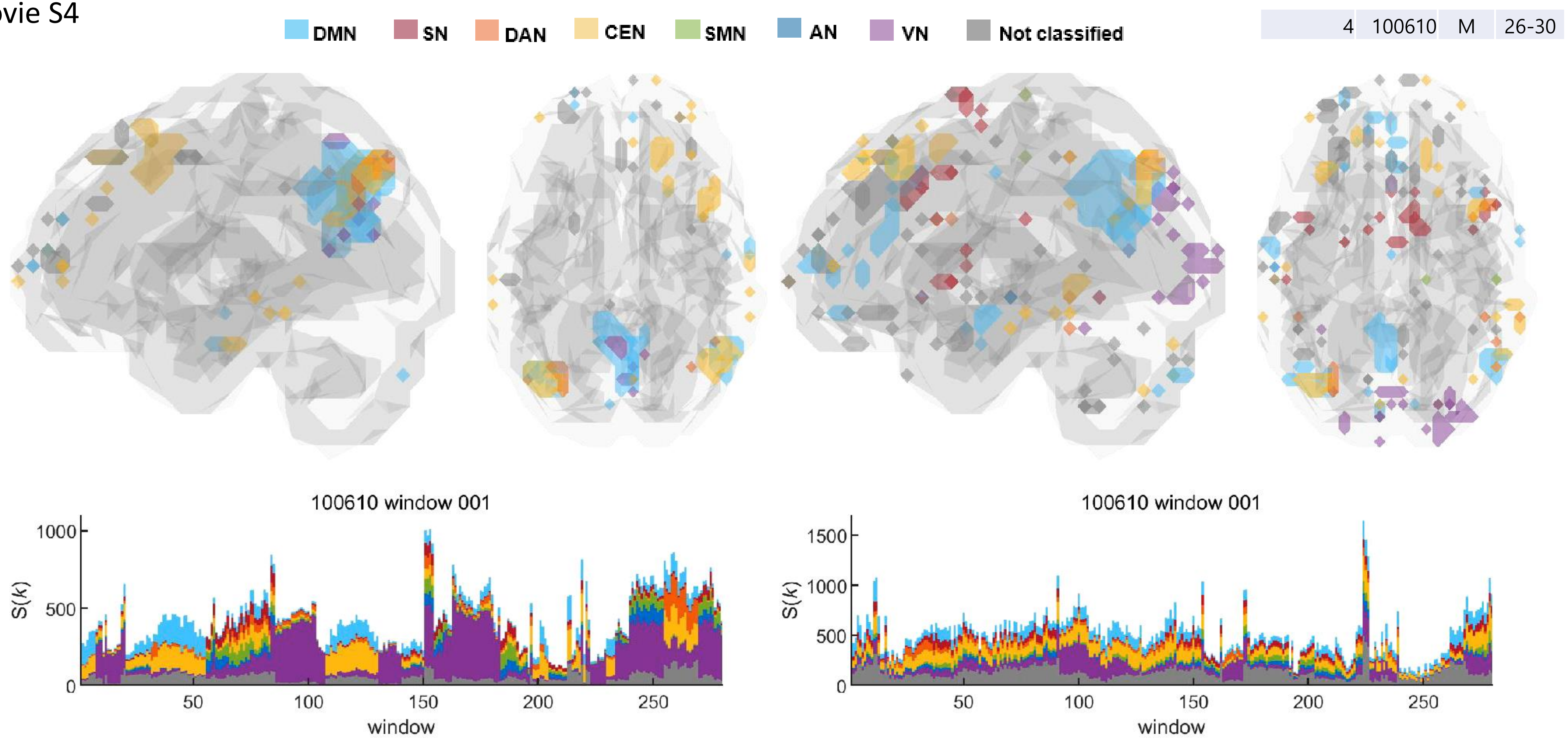


Positive correlation

Negative correlation

### Movie S5

| | | | |
|---|---|---|---|
| 23 | 104012 | F | 26-30 |

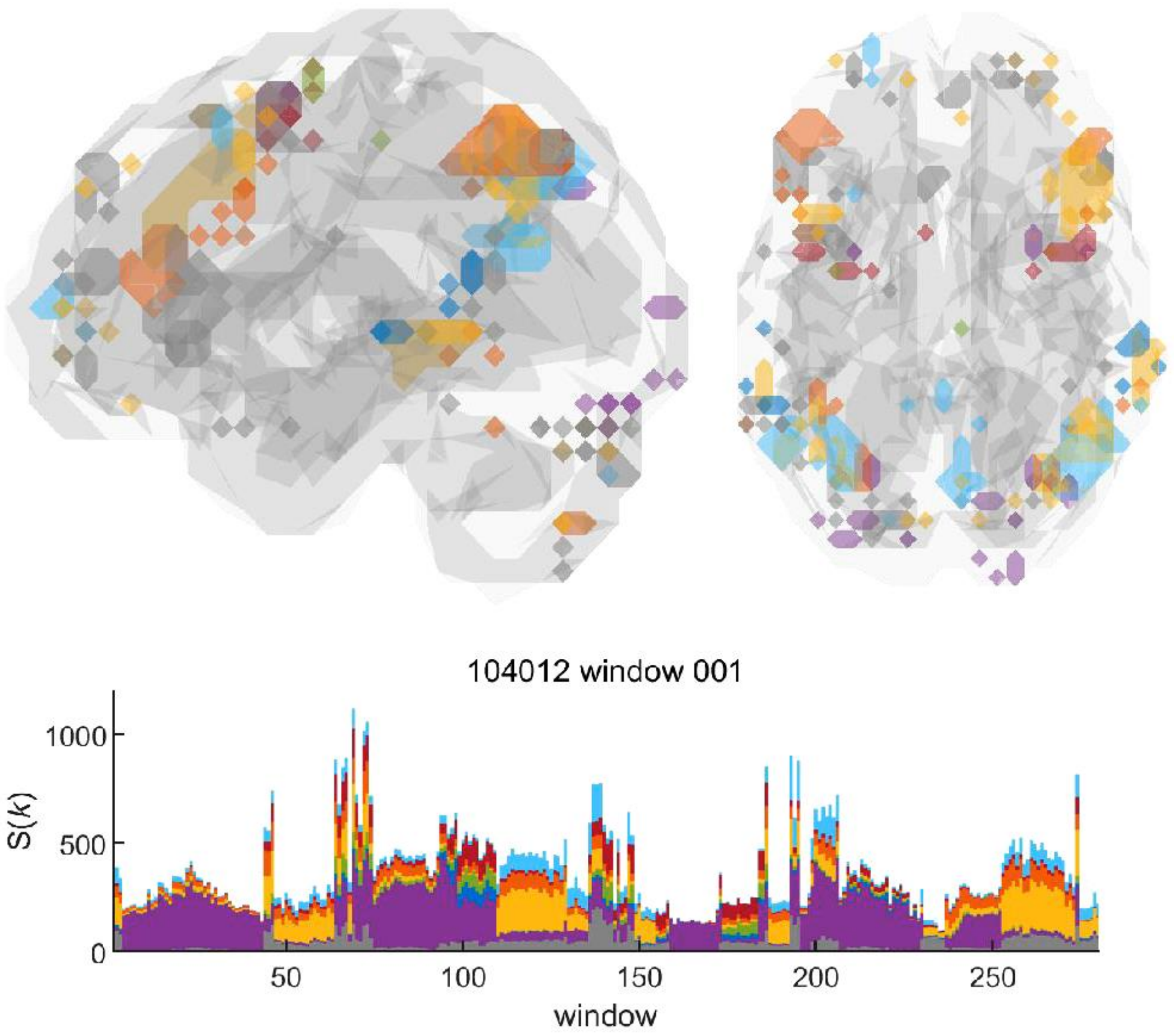


Positive correlation

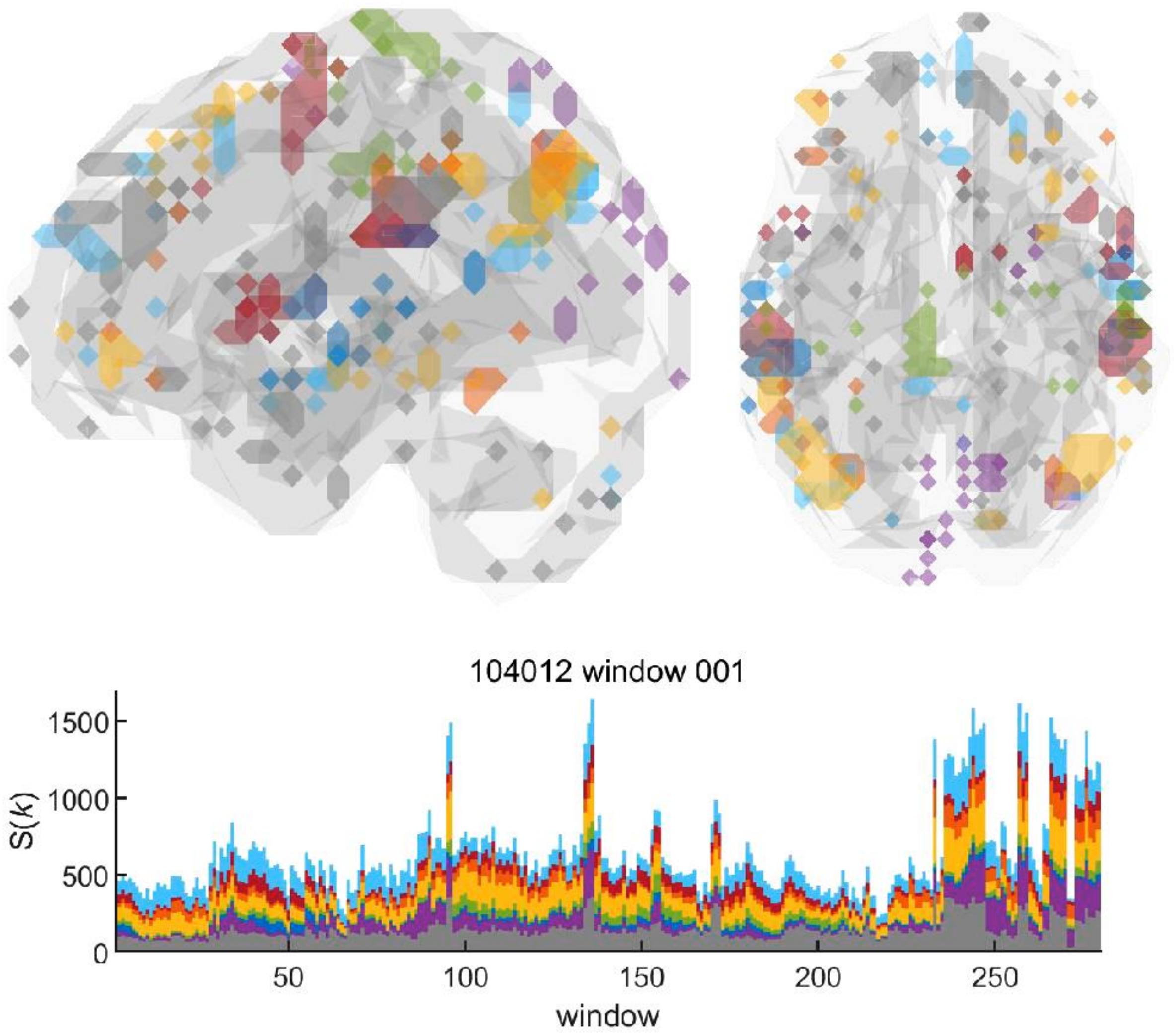


Negative correlation

Movie S6

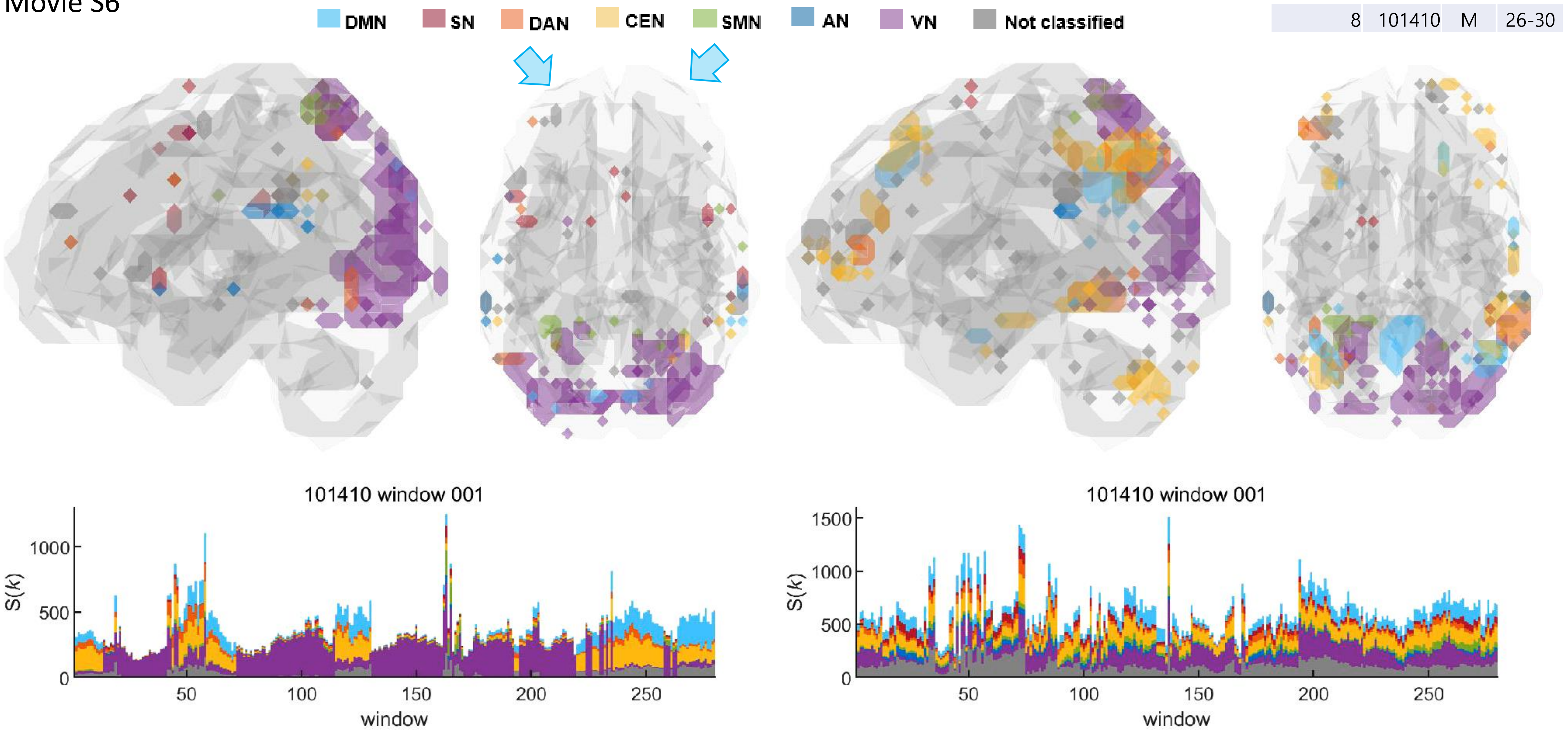


**Positive correlation**

**Negative correlation**

Movie S7

A

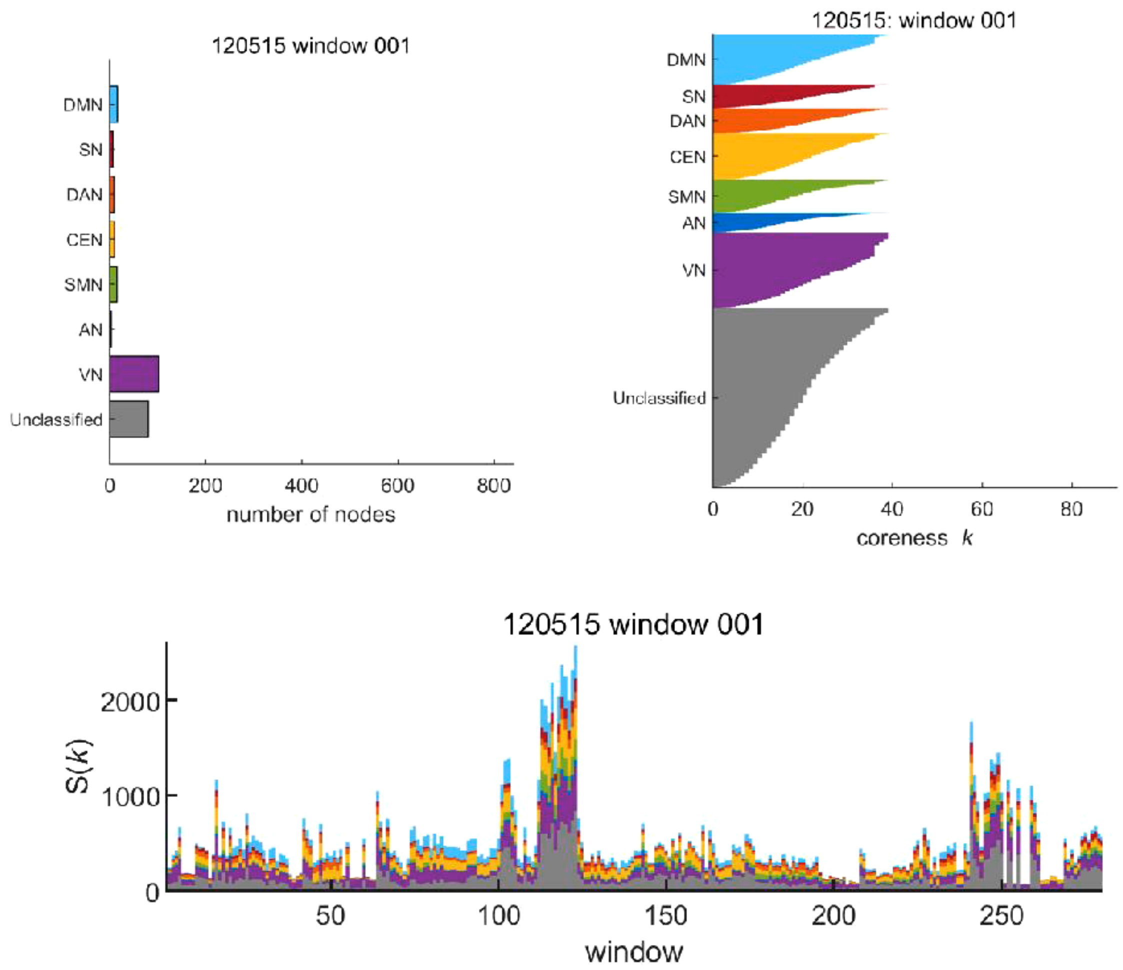


Negative correlation in a case

B

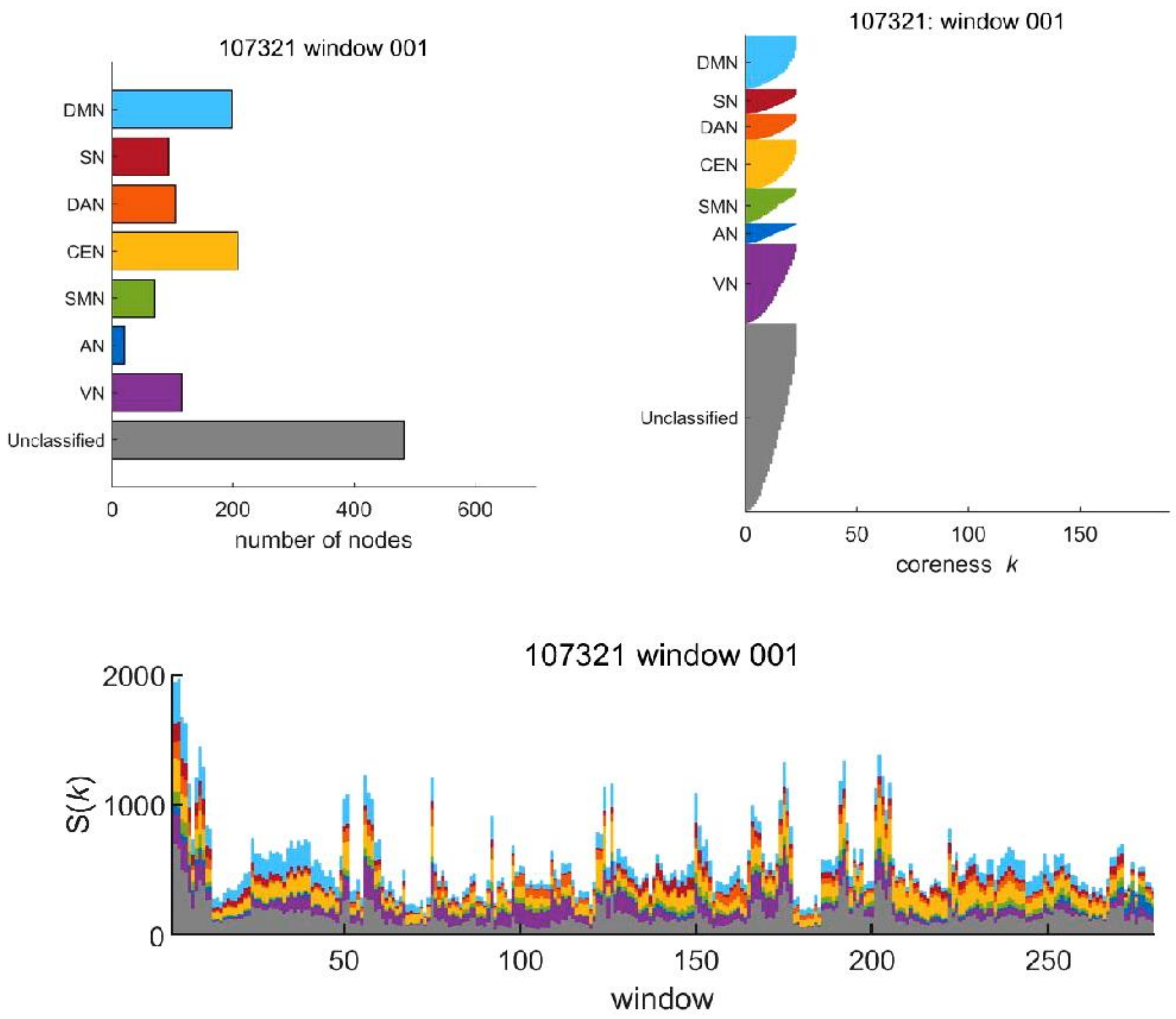


Negative correlation in another case

### Movie S8

#### Static (DMN/CEN) vs. Dynamic

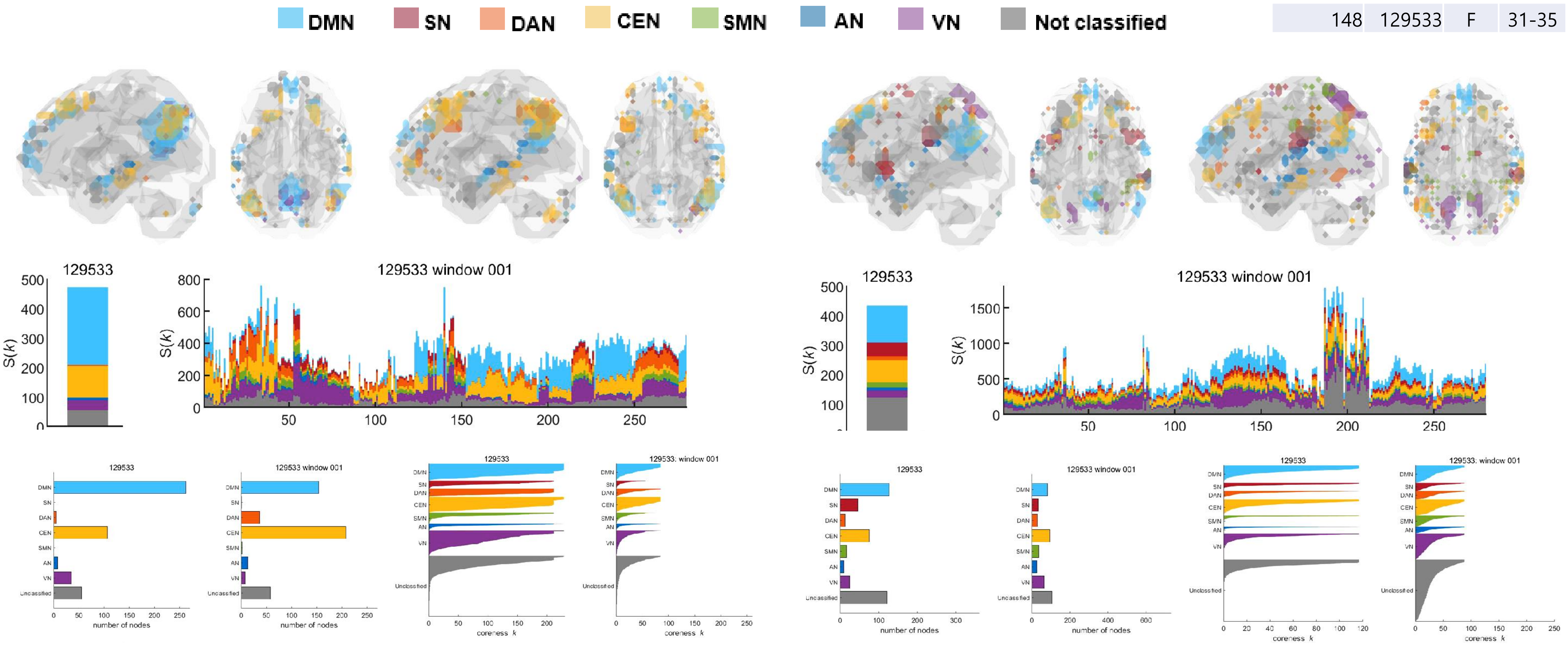


Positive Correlation

Negative Correlation

Movie S9

Static (VN) vs. Dynamic

| 131 | 126325 | F | 26-30 |
| --- | --- | --- | --- |

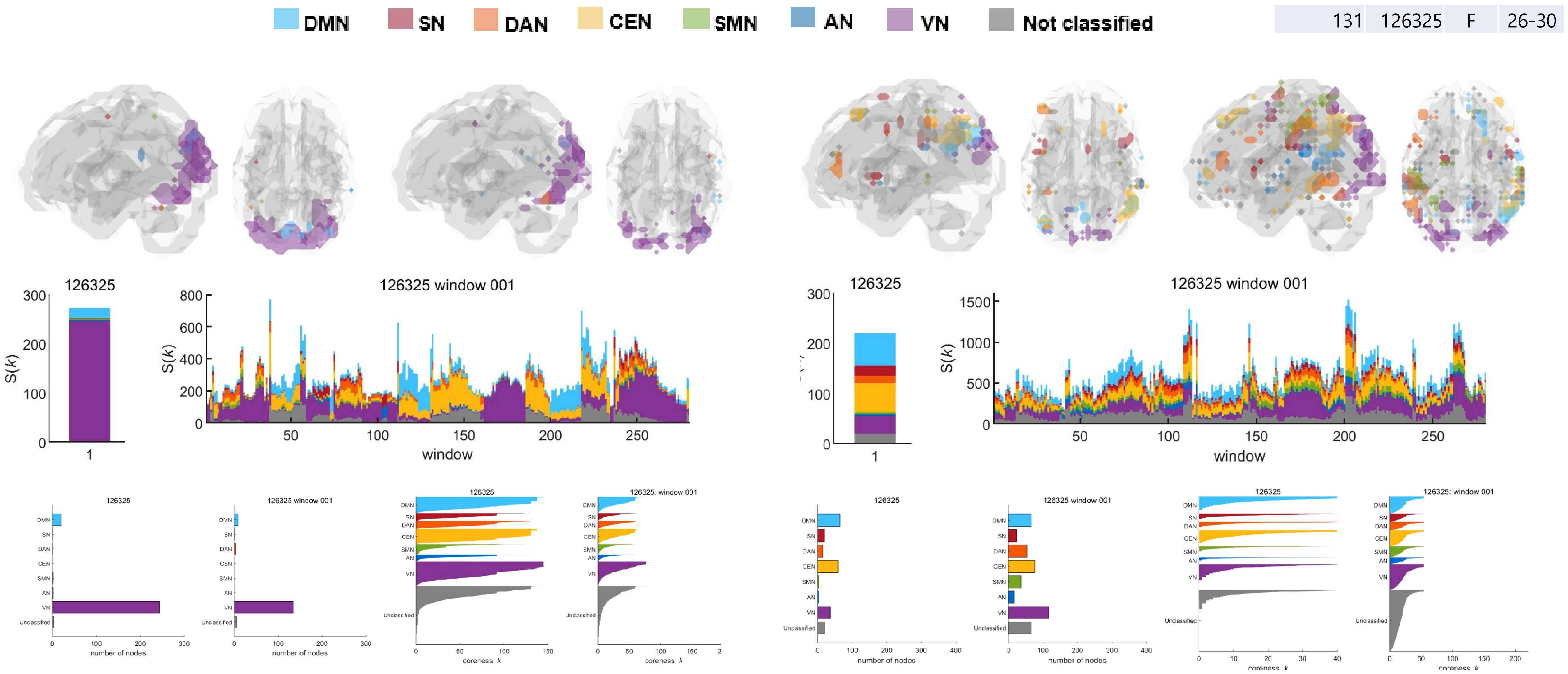


Positive Correlation

Negative Correlation

Movie S10

#### Static (Distributed) vs. Dynamic

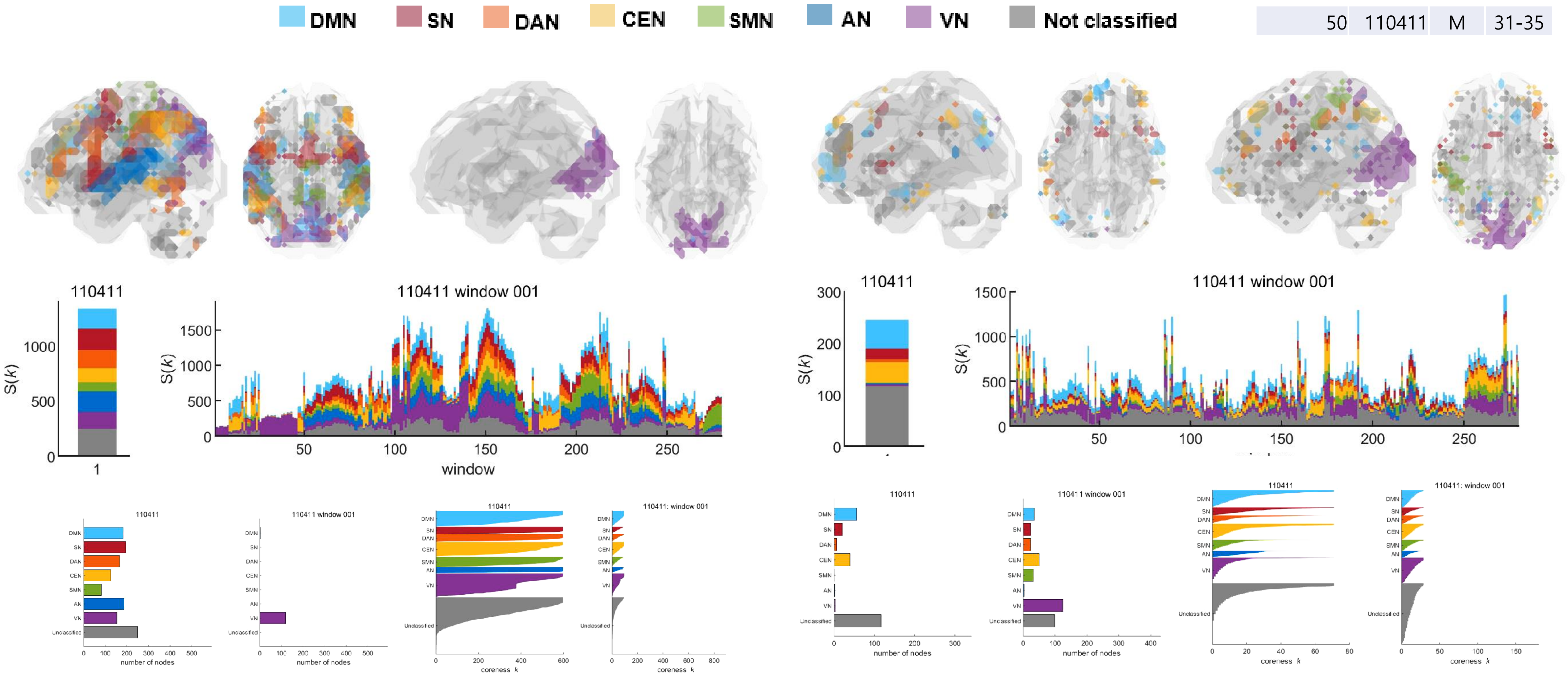


Positive Correlation

Negative Correlation

Movie S11

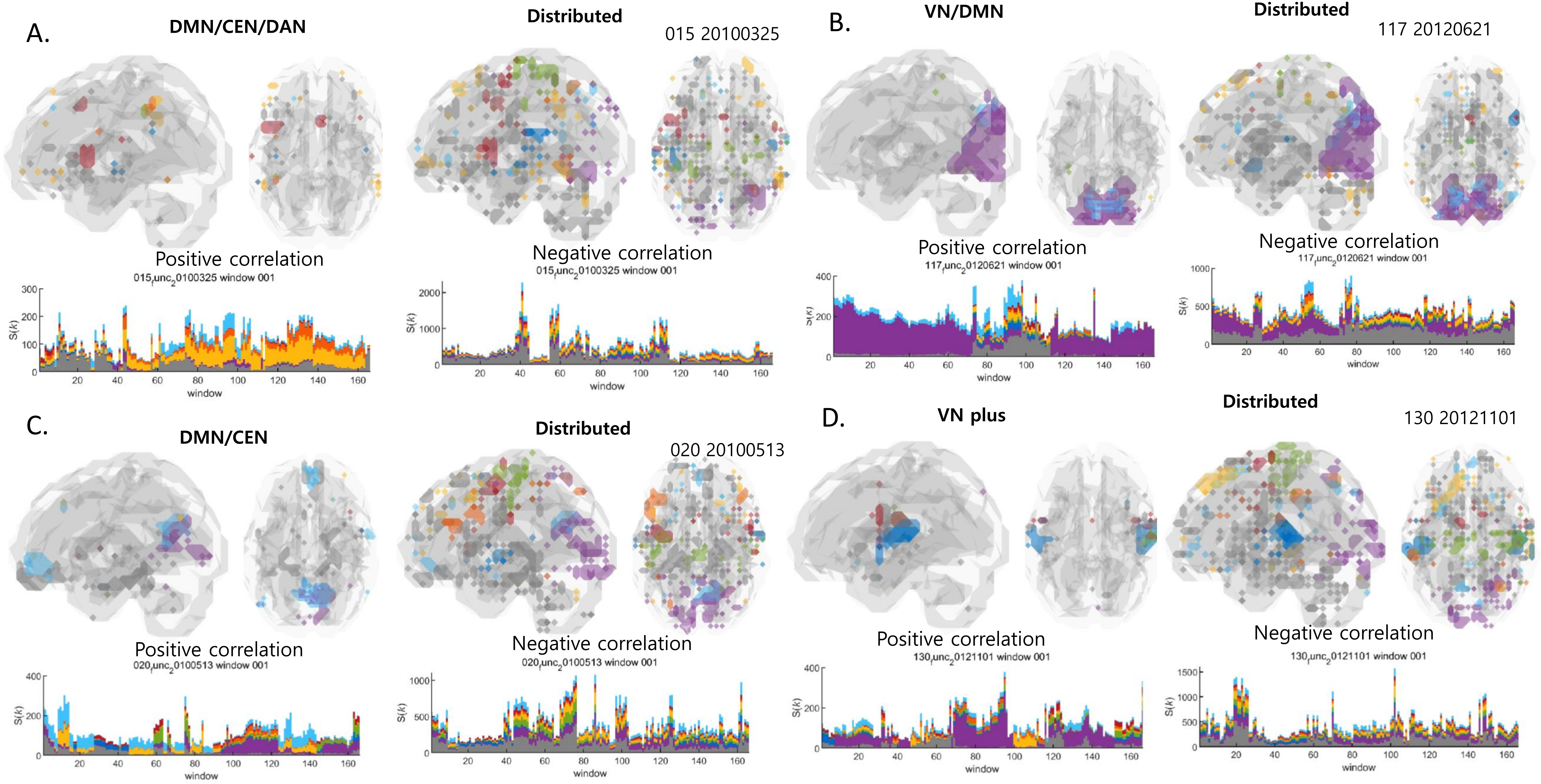

A.
DMN/CEN/DAN
Distributed
015 20100325
Positive correlation
Negative correlation
B.
VN/DMN
Distributed
117 20120621
Positive correlation
Negative correlation
C.
DMN/CEN
Distributed
020 20100513
Positive correlation
Negative correlation
D.
VN plus
Distributed
130 20121101
Positive correlation
Negative correlation
S(k)
window

Movie S12

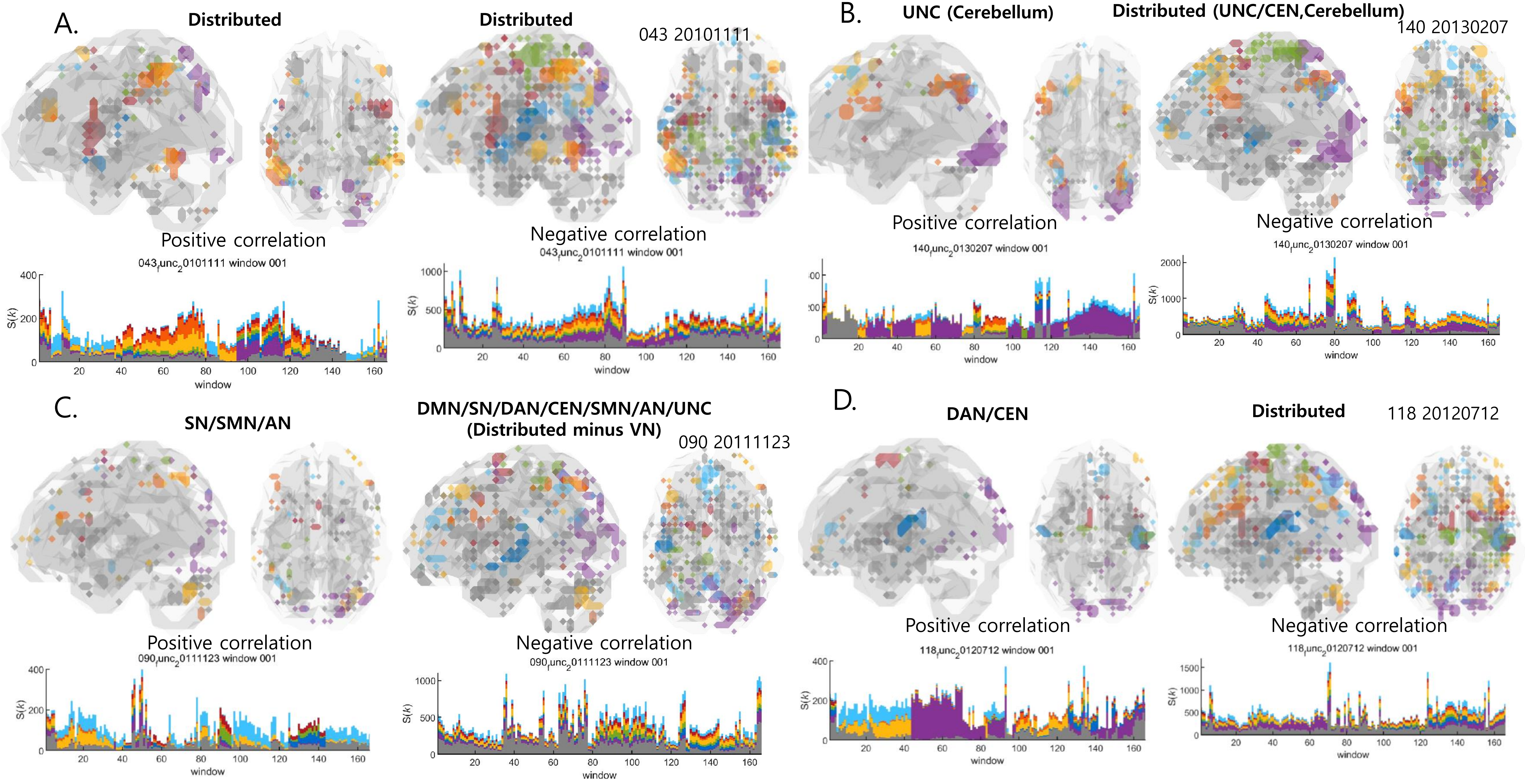
A.
Distributed
Distributed
043 20101111
Positive correlation
Negative correlation
B.
UNC (Cerebellum)
Distributed (UNC/CEN,Cerebellum)
140 20130207
Positive correlation
Negative correlation
C.
SN/SMN/AN
DMN/SN/DAN/CEN/SMN/AN/UNC
(Distributed minus VN)
090 20111123
Positive correlation
Negative correlation
D.
DAN/CEN
Distributed
118 20120712
Positive correlation
Negative correlation
S(k)
window
